# msqrob2PTM: differential abundance and differential usage analysis of MS-based proteomics data at the post-translational modification and peptidoform level

**DOI:** 10.1101/2023.07.05.547780

**Authors:** Nina Demeulemeester, Marie Gébelin, Lucas Caldi Gomes, Paul Lingor, Christine Carapito, Lennart Martens, Lieven Clement

## Abstract

In the era of open-modification search engines, more post-translational modifications than ever can be detected by LC-MS/MS-based proteomics. This development can switch proteomics research into a higher gear, as PTMs are key in many cellular pathways important in cell proliferation, migration, metastasis and ageing. However, despite these advances in modification identification, statistical methods for PTM-level quantification and differential analysis have yet to catch up. This absence can partly be explained by the inherently low abundance of many PTMs and the confounding of PTM intensities with its parent protein abundance.

Therefore, we have developed msqrob2PTM, a new workflow in the msqrob2 universe capable of differential abundance analysis at the PTM, and at the peptidoform level. The latter is important for validating PTMs found as significantly differential. Indeed, as our method can deal with multiple PTMs per peptidoform, there is a possibility that significant PTMs stem from one significant peptidoform carrying another PTM, hinting that it might be the other PTM driving the perceived differential abundance.

Our workflows can flag both Differential Peptidoform (PTM) Abundance (DPA) and Differential Peptidoform (PTM) Usage (DPU). This enables a distinction between direct assessment of differential abundance of peptidoforms (DPA) and differences in the relative usage of peptidoforms corrected for corresponding protein abundances (DPU). For DPA, we directly model the log2-transformed peptidoform (PTM) intensities, while for DPU, we correct for parent protein abundance by an intermediate normalisation step which calculates the log2-ratio of the peptidoform (PTM) intensities to their summarized parent protein intensities.

We demonstrated the utility and performance of msqrob2PTM by applying it to datasets with known ground truth, as well as to biological PTM-rich datasets. Our results show that msqrob2PTM is on par with, or surpassing the performance of, the current state-of-the-art methods. Moreover, msqrob2PTM is currently unique in providing output at the peptidoform level.

## Introduction

Mass-spectrometry-based proteomics allows the identification and quantification of a myriad of posttranslational modifications (PTMs) which reveal additional complexity and diversity of the proteome. Indeed, PTMs greatly extend the number of different forms of a protein, i.e., proteoforms, that can be found. More importantly, these PTMs can impact protein functions(1–4) and are linked to a variety of diseases and developmental disorders(5–8). Aberrant PTM status can cause a number of detrimental effects ranging from the alteration of protein folding to the dysregulation of cell signalling. It is thus of great importance to study these PTMs in detail, not only through their correct identification but also by their correct quantification and subsequent statistical analysis.

In recent years, there has been a significant improvement in the identification of PTMs with the advent of open-modification search engines such as MsFragger(9), Open-pFind(10) and ionbot(11). Yet, bespoke statistical methodologies for differential PTM analysis are lacking. To our knowledge, the only dedicated tool released at the time of writing is MSstatsPTM(12). This can be partly attributed to the complexity of PTM-rich data. Peptides can contain multiple PTM sites, sites are not always modified and modified peptides are usually harder to detect than their non-modified counterparts(4). This means that enrichment methods are most often needed for sufficient detection, which increases technical variability and experimental complexity, time and cost, which in turn leads to less available replicates(13, 14). As a result, PTM-rich data are characterised by a high amount of missingness and variability, complicating statistical analysis.

Moreover, the parent proteins on which the PTMs occur can also change in abundance regardless of the PTM. Any changes in abundance of a PTM are then confounded with changes in protein abundance(15). It is therefore crucial that any proposed statistical methodology for PTMs can take this into account.

Here, we introduce the concept of differential PTM abundance (DPA) and differential PTM usage (DPU) to enable a clear distinction between directly assessing differential abundance of PTMs (DPA) on the one hand, and differences in relative PTM abundance upon correction for the overall protein abundance (DPU), on the other hand.

In the current state-of-the-art, MSstatsPTM, DPU is achieved through an adjustment based on the model estimates of a separate PTM model as well as a protein model. We argue that this approach is suboptimal as it fails to leverage the inherent correlation between the parent protein and PTMs or peptidoforms, i.e., a specific peptide with its corresponding modifications. Additionally, the separate modelling and adjustment process in MSstatsPTM can artificially amplify small differences. This phenomenon is demonstrated in figure 1. Here, we can see a PTM for which the PTM intensities closely mimic the protein intensities across the samples. Although no significant differences are observed at the PTM or protein level when comparing the "Combo" and "Ctrl" conditions in the dataset, the adjustment inflates the difference, causing MSstatsPTM to return a significant PTM. Hence, in msqrob2PTM we employ a different normalisation strategy that directly accounts for this correlation between peptidoform and protein.

**Figure 1:**
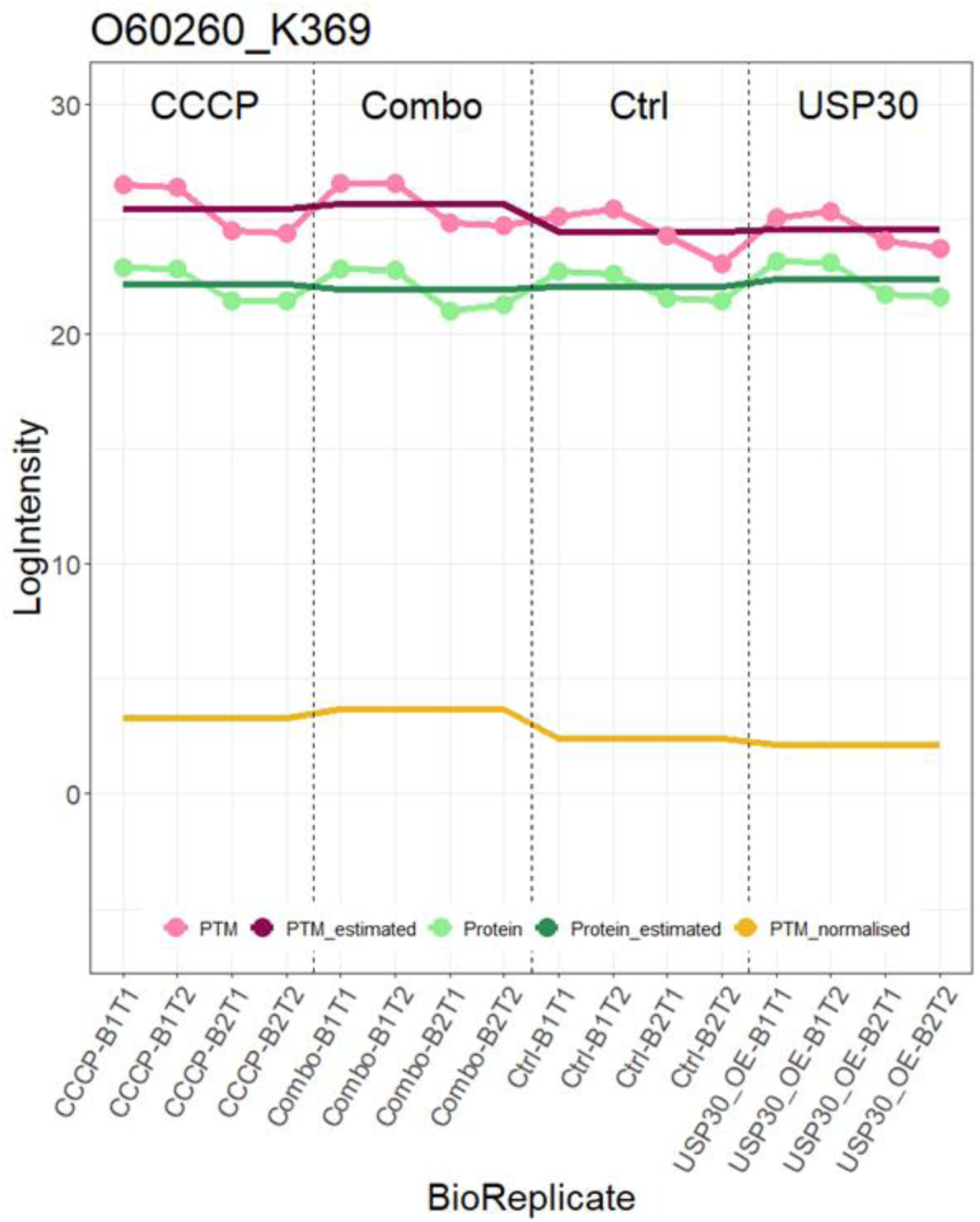
line plot displaying the PTM log_2_ intensity values (pink dotted line) and log_2_ intensity values of its parent protein (light green dotted line) in each sample. MSstatsPTM first fits a model to the PTM (dark pink line) and to the protein intensities (dark green line) to estimate the average intensity in each condition. Subsequently, the fitted average protein abundances are subtracted from the fitted average PTM intensities to obtain the average PTM abundances in each condition corrected for protein abundance (yellow line). MSstatsPTM corrected PTM abundances seem to indicate differential PTM usage. Moreover, the comparison between “Combo” vs “Ctrl” is returned by MSstatsPTM as statistically significant. This, however, appears to be an artifact of MSstatsPTM as the correction for protein abundance does not account for the link between protein and PTM intensities within samples. Indeed, when comparing “Combo” and “Ctrl” sample level intensities, the pattern at the PTM-level closely follows that of its parent protein.

Additionally, we will not limit ourselves to the analysis of the PTMs. Indeed, our method can manage the analysis of peptidoforms as well. In many studies, each distinct PTM will likely not be characterized by a myriad of peptidoforms. It is therefore possible that a significant PTM effect can be attributed to only one or two strongly significant associated peptidoforms, which may be significant for another reason, i.e. a different PTM occurring on that (those) peptidoform(s). We think it is crucial that potential users thus do not restrict their analysis to the PTM alone, but also assess the individual peptidoforms that carry the specific PTM.

We here present a statistical, R-based workflow, based on the msqrob2 R package(16), to carry out differential abundance as well as differential usage analysis at the peptidoform and PTM level. We apply this workflow to simulated datasets, a spike-in study, and to biological datasets, and use these to compare our method to MSstatsPTM. We show that our approach does not suffer from the artifacts that are introduced by uncoupling the within-sample correlation between PTM and parent protein, while maintaining good sensitivity and FDR control. The approach is freely available and can be consulted on https://github.com/statOmics/msqrob2PTMpaper

### Experimental procedures

In this section, we first introduce the msqrob2 workflow for differential peptidoform/PTM abundance and usage analysis. Next, we introduce the datasets that were used to test and validate the workflow and benchmark it to MSstatsPTM.

#### Workflow

The general workflow for the differential abundance analysis on PTM and peptidoform level was developed in R(17) (version 4.2) and is mainly based on two R packages: msqrob2 (https://www.bioconductor.org/packages/release/bioc/html/msqrob2.html, version 1.6.0) and QFeatures(18) (https://www.bioconductor.org/packages/release/bioc/html/QFeatures.html, version 1.8.0). QFeatures provides an infrastructure to store and manage mass spectrometry data across different levels (e.g. peptidoform and protein level) whilst keeping links between the levels where possible. For each preprocessing step a novel, linked assay is constructed. In this way, the original data is not overwritten, and preprocessed data can be traced back to its origin. msqrob2 is a package with updated and modernised versions of the MSqRob(16) and MSqRobSum(19) tools and builds upon the QFeatures class infrastructure. It provides a robust statistical framework for differential analysis of label-free LC-MS proteomics data to infer on differential abundance on the peptide (peptidoform) and/or protein level. Here, we add workflows that provide inference on differential abundance and usage at the PTM and peptidoform level.

We make a distinction between differential abundance and differential usage. This is the difference between directly assessing differential abundance (DA) on the one hand, and differences in relative abundance upon correction for the overall protein abundance (DU), on the other hand. Essentially, this relates to a difference in normalisation (see point 3 below).

We first provide an overview of the workflow before going over each step in detail.

1. Conversion of input data and construction of the QFeatures object
2. Pre-processing
3. Normalisation
4. Peptidoform level analysis
5. Summarisation of peptidoforms to PTM level
6. PTM level analysis
7. Results exploration plus visualisation

##### 1. Conversion of input data and construction of the QFeatures object

As input data, we require the output of a quantification algorithm (in txt or csv format) that contains all peptidoform identifications, parent protein(s) and per sample intensities. This should be in wide format: each unique peptidoform should be on one line that contains (at least) the information on its parent protein, modification (plus location), and intensities for each sample. As quantitative proteomics data can be readily transformed into this format, we have no restrictions on search engines or quantification algorithms users want to adopt.

Once the data are in the right format they are imported as a QFeatures object. Next, information on the experimental design can be added in the *colData* instance of the object.

##### 2. Pre-processing

First, the peptidoform data can be filtered. Each peptidoform should have measured intensity values in at least two samples, or else are filtered out. Intensities are log-transformed if not already the case. Of course, decoys and contaminants should be removed.

The pre-processing steps are not limited to those above, as, depending on the nature of the dataset and user knowledge, more filtering steps can be added.

##### 3. Normalisation

Distinct normalisation steps should be adopted for inferring on differential abundance and differential usage. For DA, only median centring or mean centring can be used, e.g. via the *normalise* function from the QFeatures package. DU requires an additional normalisation to correct for changes occurring in the parent protein. Indeed, changes in the overall protein abundance between conditions can trigger the associated PTM(s) to be detected as differentially abundant. To infer on PTM(s) for which the effect of the treatment differs from that of the overall protein, we first summarise the protein intensity value per sample for each unique protein, e.g., via robust regression using the *robustSummary* function in the MsCoreUtils(20) R package, and we subsequently subtract it from the intensity values corresponding to all peptidoforms derived from that protein, i.e.

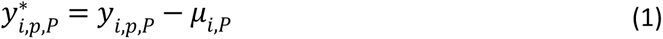

With 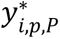 the normalised log2-transformed intensity for peptidoform *p* in sample *i* with parent protein *P*, *y_i,p,P_* the log2-transformed intensity for peptidoform *p* in sample *i* with parent protein *P* before normalisation and *μ_i,P_* the summarised intensity for protein *P* in sample *i*.

It is possible to calculate the summarised protein intensity value directly from the PTM dataset itself. However, when the experiment includes both an enriched and non-enriched (global profiling) dataset we recommend using the non-enriched dataset to calculate the summarised protein values. Of note, steps one and two should also be applied to the non-enriched data.

##### 4. Peptidoform level analysis

Before transitioning to the PTM level, it is possible to directly assess differential usage or expression on peptidoform level. The steps to take are exactly the same as step 6 below, but instead of using the PTM assay obtained in step 5, we use the normalised peptidoform assay obtained in step 3 as input to the *msqrob* function.

This allows the user to assess associated peptidoforms underlying significant PTMs of interest.

##### 5. Summarisation of peptidoforms to PTM level

For each unique PTM (i.e. unique protein – modification – location combination), we need a summarised intensity value per sample. This is done by taking a subset of the dataset with all peptidoforms containing a specific PTM and summarising all corresponding intensity values into one value per sample. When peptidoforms contain multiple PTMs, these are used multiple times. Here we apply robust regression using the *robustSummary* function in the MsCoreUtils(20) R package by default to summarise the peptidoform level data at the PTM-level. In this way, we obtain an intensity assay on the PTM level. This assay can then be added to the existing QFeatures object.

##### 6. PTM level analysis

We use the functionalities of the msqrob2 package for this step. Msqrob2 (16, 19, 21, 22) provides a robust linear (mixed) model framework for assessing differential abundance in proteomics experiments. To assess differential abundance on the protein level, the workflows can start from raw peptide intensities or summarised protein abundance values. The model parameter estimates can be stabilized by ridge regression, empirical Bayes variance estimation and robust M-estimation. Here we assess differential abundance on the PTM level by first summarising peptidoform expression values (step 5).

When one predictor (e.g. *condition*) is present in the dataset, we perform an msqrob analysis on PTM intensities with the following model:

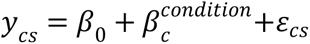

With *y*_*cs*_ the summarised log2-transformed PTM intensity in sample *s* of condition *c*, *β*_0_ the intercept, and 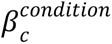, the effect of a condition *c*. The error term ε_*cs*_ is assumed to be normally distributed with mean 0 and variance σ2.

When multiple predictors are present, the model can be expanded as needed, with the additional possibility of using mixed models. The user needs to specify the model formula themselves using lm or lme4(23) R syntax.

The contrast matrix for contrasts of interest can be specified via the *makeContrast* function present in msqrob2, which are subsequently assessed using the *hypothesisTest* function. By default, the results of the latter function are corrected for multiple testing using the Benjamini-Hochberg false discovery rate (FDR) method.

The model results are stored in the existing QFeatures object together with the raw data and the pre-processed data.

##### 7. Results exploration plus visualisation

The abovementioned model results contain a significance table with (adjusted) p-values, log fold changes, standard errors, degrees of freedom and test statistics.

Different visualisations can easily be made based on this table and the links to the underlying intensity data in the QFeatures object, such as volcano plots, heatmaps and line plots at the peptidoform, PTM and/or protein level.

#### Data

Our novel msqrob2 workflow is tested and benchmarked to MSstatsPTM using two computer simulations developed by the MSstatsPTM team, the spike-in dataset from the MSstatsPTM paper, and data from two real experiments.

The computer simulations were specifically developed for testing differential PTM workflows, and also allowed us to directly compare our method to MSstatsPTM. The first simulation produced “perfect” datasets with no missing values and many modified features per PTM, while the second simulation incorporated missing values and limited modified features, producing more lifelike datasets.

The spike-in dataset consists of fifty human ubiquitinated peptides that were spiked into four background mixtures in known amounts. Hence, the true log-fold changes and identity of the truly changing PTMs are known.

The biological case studies consist of a biological, label-free LC-MS/MS ubiquitination experiment by Cunningham *et al*. to study the role of USP30, a deubiquitylase enzyme, in mitophagy regulation(24); and a label-free phosphorylation experiment with a two-factor design where both the total proteome and phosphoproteome were measured.

More details on each dataset are given below.

### Computer simulations

We used the two computer simulations from the MSstatsPTM team that were found on https://github.com/devonjkohler/MSstatsPTM_simulations/tree/main/data (simulation1_data.rda and simulation2_data.rda). The first simulation consists of data without any missing values, while in the second simulation, missing data is introduced. For each simulation, 24 datasets were created with different experimental designs and intensity variance. In each dataset, 1000 PTMs were simulated.

Half of the PTMs were simulated to have a fold change between conditions. However, of the half with differential fold changes on the PTM level, 250 could be confounded with differential fold changes of the parent protein. For further details on the creation of the datasets, we refer to the MSstatsPTM paper(12) and to their GitHub page.

Both simulations contain an enriched PTM dataset as well as its non-enriched protein counterpart. From each of the 24 datasets, the FeatureLevelData was extracted from the PTM and the protein dataset. These two datasets were then used as input to the workflow and all seven steps were followed. The protein dataset was used for the normalisation step.

Because it is known which PTMs are differentially abundant and/or differentially used, we can readily evaluate the performance of a method in terms of the false positive rate (fpr), sensitivity, specificity, precision and accuracy, and true positive rate (tpr) - false discovery proportion (fdp) plots. Note, that tpr is the fraction of the truly differentially abundant PTMs picked up by the method and fdp is the fraction of false positives in the total number of PTMs flagged as differentially abundant. On the tpr-fdp plot we also indicate the observed fdp at 5% FDR cut-off, which is expected to be close to 5%.

We compared our results with the results obtained with the MSstatsPTM method, on their GitHub page https://github.com/devonjkohler/MSstatsPTM_simulations/tree/main/data

(adjusted_models_sim1.rda and adjusted_models_sim2.rda) and included these in the tpr-fdp plots.

### Spike-in dataset

The MSstatsPTM team also developed a biological spike-in dataset with known ground truth to test their approach. Fifty human ubiquitinated peptides were spiked into four background mixtures consisting of human and E.coli proteins in different amounts. These four mixtures represent four different conditions and for each, two replicates were created. An overview of the experimental design can be seen in figure 2. Because the amount of spiked-in peptides is known, the true log-fold changes between the conditions is known and it is possible to assess whether the method can pick up these fold changes, and if these fold changes differ from the fold change of the corresponding protein in the background. Note, however, that as opposed to real experiments, the ubiquitinated peptides in the spike-in study are not correlated to their corresponding protein in the background. Further technical details can be found in the MSstatsPTM paper (12). The dataset can be found on MassIVE: <u>MSV000088971</u>.The true log fold changes (before and after protein adjustment) are depicted in table 1.

**Figure 2:**
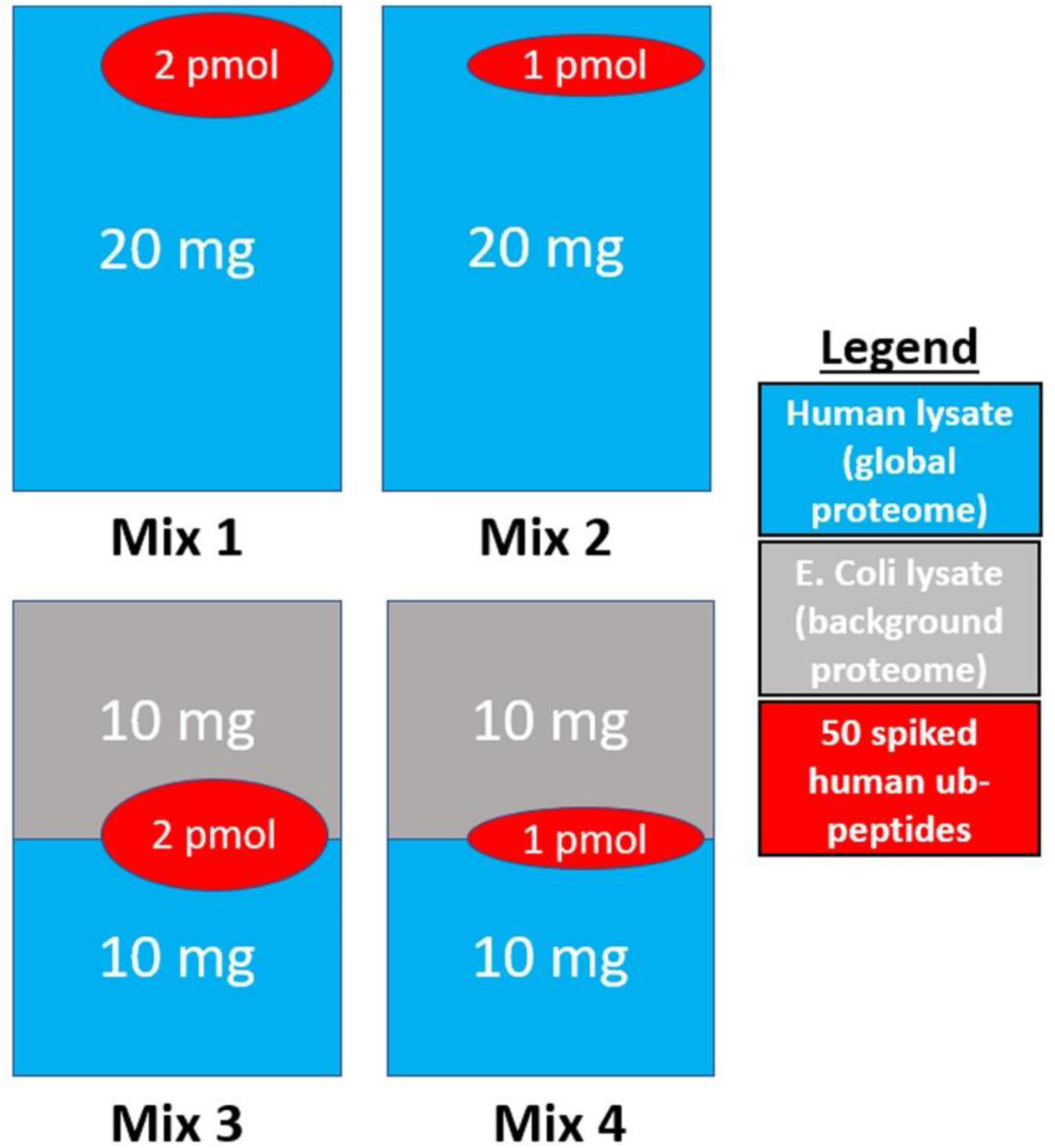
Experimental design of the spike-in dataset. Fifty human heavy labelled KGG motif peptides were spiked into four background mixtures in different amounts. Mixes 3 and 4 consist of a mix of E. Coli and human proteins. Only the human proteome was utilised as the global proteome. Figure adapted from (12)

**Table 1:** True log2 fold changes of the spike-in peptides in the different comparisons between the mixtures.

| Comparison | True log2 FC without adjustment | True log2 FC after adjustment |
| --- | --- | --- |
| mix2 vs mix1 | -1 | -1 |
| mix3 vs mix1 | 0 | 1 |
| mix4 vs mix1 | -1 | 0 |
| mix3 vs mix2 | 1 | 2 |
| mix4 vs mix2 | 0 | 1 |
| mix4 vs mix3 | -1 | -1 |

As input to our workflow, we used the MSstatsPTM_Summarized.rda object provided on MassIVE. In the FeatureLevel data part of the object, the spiked-in peptides were not annotated and were irretrievable because the heavy peptides can also be present as their light counterparts. However, they were annotated in the ProteinLevel part. Hence, we could not use the low-level data, and had to start from the data that had already been pre-processed and summarised to PTM (for the PTM dataset) and Protein level (for the global profiling dataset) by MSstatsPTM, thus omitting step 4 and 5 from the workflow. We therefore could not assess our entire workflow based on these data, and moreover do not know which preprocessing steps were conducted.

We employed various methods to analyse this dataset. Our primary approach was the msqrob2PTM workflow as described in the workflow section, as well as the normal MSstatsPTM workflow. We also assessed the differential abundance of the PTMs with the standard msqrob2 workflow: *DPA-nonNorm*, which does no normalisation and hence skips step 3 of the standard workflow entirely and *DPA*, which only applies median centring in step 3.

Because we know the ground truth of this dataset, we can again use the same metrics to assess the performance. Here, we also make ROC (tpr-fpr) curves. Furthermore, the log fold changes estimated by msqrob2PTM and MSstatsPTM were used to generate boxplots showing the observed and expected FCs for each mixture. For MSstatsPTM, the log fold changes were derived from the MSstatsPTM_Model.rda object and Spike-in_Vizualization.Rmd contained R code for the boxplots. Both these files were found on <u>MassIVE RMSV000000669</u>.

### Ubiquitination dataset

Details of the experimental set-up can be found in reference (24). The dataset itself is available on MassIVE(25) as <u>MSV000078977</u>.

The dataset consists of four conditions: carbonyl cyanide 3-chlorophenylhydrazone (CCCP) treatment, USP30 overexpression (USP30-OE), a combination of both (Combo), and a control group. Per condition, two biological replicates with two technical replicates each were generated. All pairwise comparisons were tested using msqrob2PTM.

This dataset has also been used in the MSstatsPTM paper, hence, we can compare our results to theirs for a biological case with unknown ground truth. As input to the msqrob2PTM workflow, we used the usp30_input_data.rda object found in the MassIVE MSstatsPTM analysis container RMSV000000358, which was also used as input to the MSstatsPTM workflow. This ensures compatibility of the results with those in the MSstatsPTM paper. In this container, the analysis file MSstatsPTM_USP30_Analysis.R can also be found, which was used for the MSstatsPTM results.

All steps of the workflow were followed as described above. The normalisation step made use of the available PTM dataset, given the lack of a non-enriched counterpart. Because each condition consists of two biological replicates which in turn consists of two technical replicates, we used the *msqrob* function with a mixed model as input.

The results of both analyses were used to generate line plots with input as well as our normalised PTM level-data and the estimated effects for each condition. The detailed model results in the MSstatsPTM model object allowed us to inspect the model output for each PTM and protein as well as those for PTM upon correction for protein.

### Phospho dataset

The human phosphorylation datasets consist of 47 samples from condition A and 43 from condition B. Two aliquots were processed for each sample: one dedicated to total proteome analysis, and the other one to the phosphoproteome analysis. The main sample preparation steps were identical for proteomics and phosphoproteomics apart from the additional phosphopeptide enrichment step. Briefly, MeOH precipitation was performed on all samples and protein pellets were resuspended with 0.1% RapiGest^TM^ surfactant (Waters). Either 20 µg (proteomics) or 100 µg (phosphoproteomics) of samples were subjected to overnight trypsin/lysC (Mass Spec Grade mix, Promega, Madison, USA) digestion at 37°C with an enzyme:protein ratio of 1:25. Peptide samples were then incubated for 45 minutes at 37°C and centrifuged to remove RapiGest.

For total proteome analysis, collected supernatants were loaded on an AssayMAP Bravo (Agilent) for automated peptide clean-up using C18 cartridges. Desalted peptides were injected on a nanoAcquity UltraPerformance LC^®^ (UPLC^®^) device (Waters Corporation, Milford, MA) coupled to a Q-Exactive Plus mass spectrometer (Thermo Fisher Scientific, Waltham, MA) and analysed using Data Dependent Acquisition (DDA).

For phosphoproteomics, collected supernatants were loaded on an AssayMAP Bravo (Agilent) for automated Fe(III)-NTA phosphopeptides enrichment. Enriched samples were then analysed on a nanoAcquity UPLC devise (Waters) coupled to a Q-Exactive HF-X mass spectrometer (Thermo Scientific, Bremen, Germany) using DDA.

Generated raw data files were searched against a database containing all human entries extracted from UniProtKB-SwissProt (25/08/2021, 20 339 entries) using MaxQuant (v.1.6.17). The minimal peptide length required was seven amino acids and a maximum of one missed cleavage was allowed. For proteomics data, methionine oxidation and acetylation of proteins’ N-termini were set as variable modifications and cysteine carbamidomethylation as a fixed modification. For phosphoproteomics data, serine, threonine and tyrosine phosphorylations were added as variable modifications. For protein quantification, the “match between runs” option was enabled. The maximum false discovery rate was set to 1% at peptide and protein levels with the use of a decoy strategy. Intensities were extracted from the Evidence.txt file to perform the following statistical analysis. All seven steps of the workflow were performed. The dataset can be found on PRIDE (PXD043476).

### Mock analyses

For the phospho dataset a mock analysis was included, that is an analysis where we only take one treatment arm of the data, so none of the PTMs (peptidoforms) are expected to be differential. We then assign the samples at random to a mock treatment with two levels and assess differential usage between the two conditions (mock vs control). In this way, correct control of the type I error by the statistical method can be assessed. Indeed, every PTM that is called as differentially abundant is a false positive. Hence, we expect the method to return uniform p-values.

From the phospho dataset, only the samples from factor 1 condition B, and factor 2 condition y were withheld, i.e. 26 samples. Upon step 4, 13 out of the 26 samples were randomly assigned to condition “mock”, the other 13 were assigned as condition “control”. Step 5 was then carried out by testing for a condition effect and the calculated p-values were retained. The randomisation to the mock treatment and step 5 in the analysis was repeated 5 times and a histogram was made for the p-values for each mock simulation.

This mock analysis was done for different workflows: we assessed the effect of using robust regression in the modelling step, the use a non-enriched counterpart for normalisation and normalisation based on the enriched dataset, itself. Moreover, we conducted the analysis both on peptidoform as well as PTM-level.

## Results

The performance of our novel PTM and peptidoform msqrob2 based workflows will be compared to MSstatsPTM based on computer simulations, the spike-in dataset, the ubiquitination and phospho datasets.

### Computer simulations

#### PTM-level

We first evaluated our method using the two computer simulations mentioned above. The first simulation consisted of 24 “perfect” datasets with no missing data and ten distinct peptidoforms carrying a specific PTM. Half of the datasets were simulated with a standard deviation of the difference in log-intensities between modified and unmodified peptidoforms of 0.2, the other half had a standard deviation of 0.3. The datasets differ in the number of replicates as well as in the number of conditions.

Figure 3 shows the true positive rate (tpr, the fraction of the truly differentially abundant PTMs picked up by the method) - false discovery proportion (fdp, the fraction of false positives in the total number of PTMs flagged as differentially abundant) curve for simulation 1 for all 24 datasets. As expected, both msqrob2PTM and MSstatsPTM perform better in datasets with lower variability and/or a higher number of replicates. Indeed, the true positive rate or sensitivity is higher for the same level of the false discovery proportion when the number of repeats increases while keeping the sd fixed (or when reducing the sd while keeping the number of repeats fixed). msqrob2PTM (solid line) clearly outperforms MSstatsPTM in all datasets (dotted line). Furthermore, MSstatsPTM in particular seems to have issues when the number of replicates is low. Indeed, in four out of six datasets with two replicates, the dotted line immediately veers right instead of up, indicating that non-DU PTMs are returned among the most significant features. This particularly affects datasets with higher variation (sd 0.3). msqrob2PTM, however, does not suffer from a poor ranking of the PTMs for these four datasets and is still able to report (a few) true positive results at the 5% FDR level. Moreover, the fdp at the 5% FDR level for msqrob2PTM is close to 5% for most datasets, indicating a good control of false positives.

**Figure 3:**
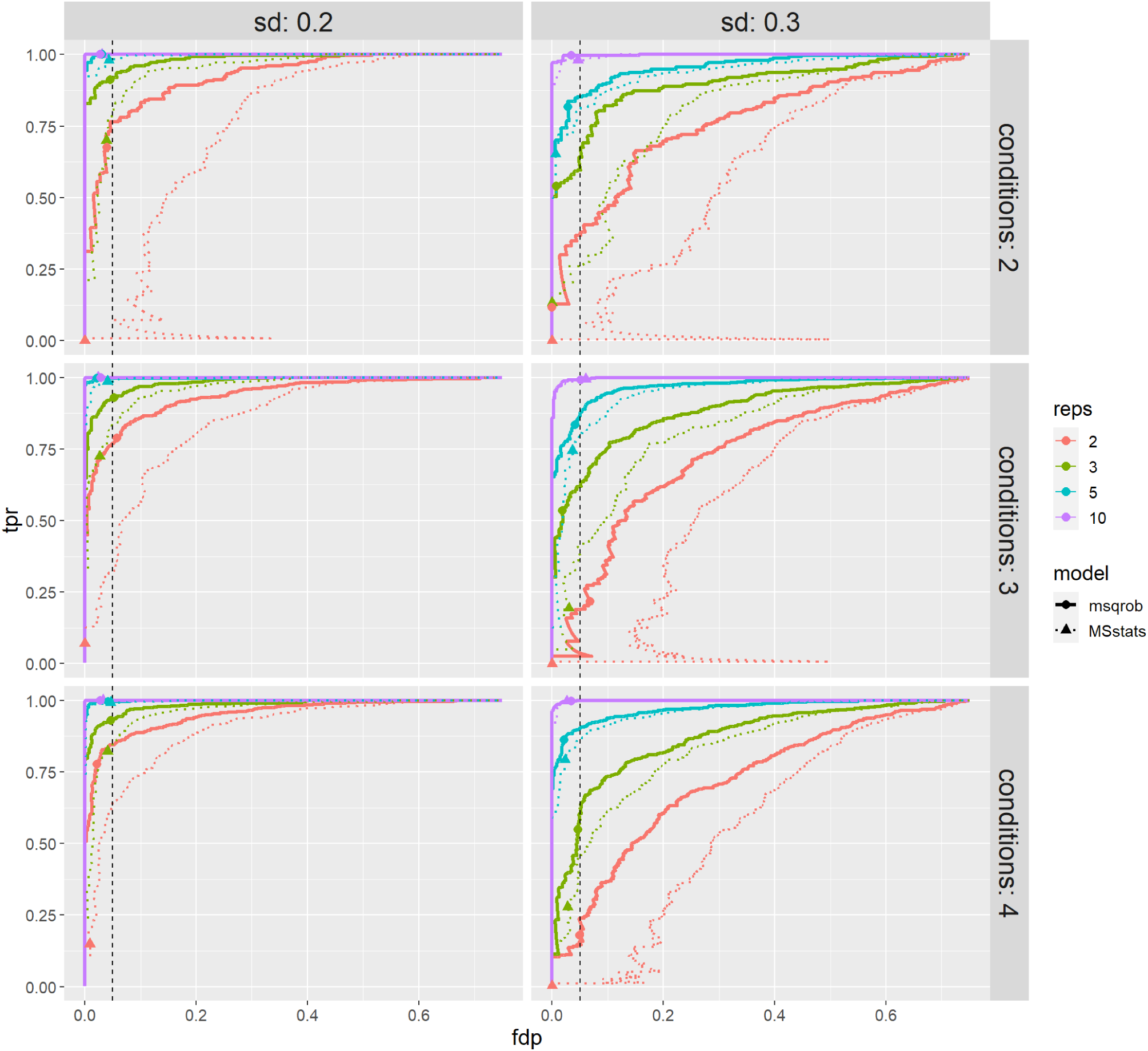
True positive rate (tpr) - false dicovery proportion (fdp) plots for datasets simulated under first scenario (no missingness). msqrob2PTM (full lines) is compared to MSstatsPTM (dotted lines). Observed fdp at a 5% FDR cut-off is denoted by dots for msqrob2PTM and by triangles for MSstatsPTM. msqrob2PTM uniformly outperforms MSstatsPTM for all datasets. Indeed, MSstatsPTM is less sensitive, i.e. its tpr-fdp curve is always below the corresponding one of msqrob2PTM.

Figure 4 shows the tpr - fdp curves for simulation 2 for all 24 datasets. As expected, the higher number of missing values induces a slight drop in performance overall. However, for the larger sample sizes the performance remains very good for msqrob2PTM. Again, msqrob2PTM uniformly outperforms MSstatsPTM and the fdp is close to 5% when adopting a 5% FDR threshold. For two datasets, we see that the far end of the tpr-fdp curve for msqrob2PTM veers straight up (two conditions, two replicates sd 0.2 and sd 0.3), which reflects msqrob2’s inability to fit the models for a number of PTMs. This happens because these PTMs have too few observations to fit the models due to the missingness introduced in this simulation scenario.

**Figure 4:**
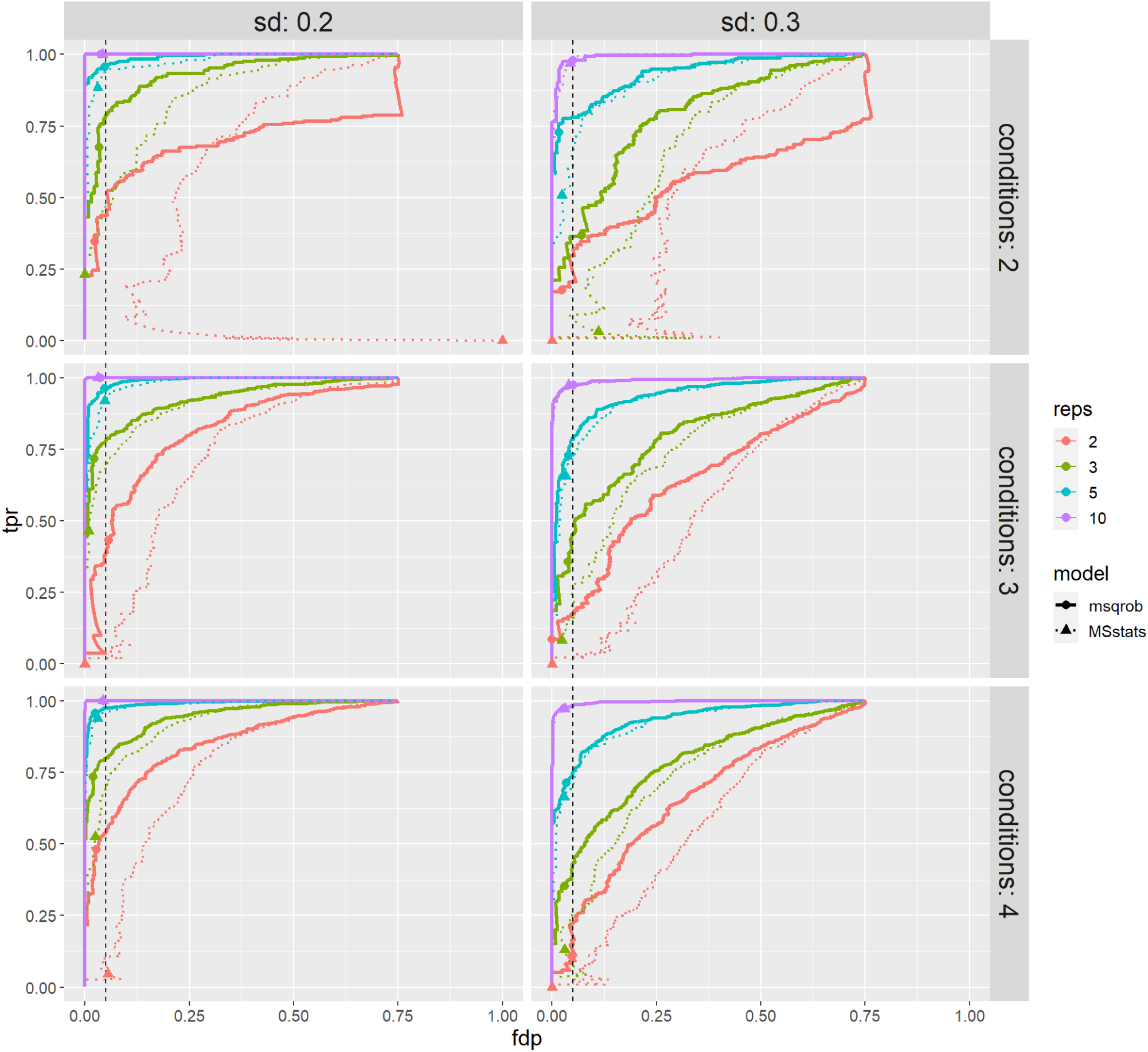
tpr-fdp plot for datasets simulated under second scenario (with missingness). msqrob2PTM (full lines) is compared to MSstatsPTM (dotted lines). Observed fdp at a 5% FDR cut-off is denoted by dots for msqrob2PTM and by triangles for MSstatsPTM. Here again, msqrob2PTM outperforms MSstatsPTM for all datasets.

For further comparison, ROC curves (true positive rate vs false positive rate) are shown in Supplementary figure 1 and 2. These plots give less weight to a few top-ranked false positives. Again, these ROC curves demonstrate superior msqrob2PTM performance.

In supplementary table 1 and 2, the performance metrics (false positive rate, sensitivity, specificity, precision and accuracy) that were reported in the MSstatsPTM paper are also given for all datasets for comparison.

#### Peptidoform level

Our msqrob2PTM workflow can also infer on differential usage at the peptidoform level, which we consider to be very important. Indeed, not all peptidoforms that carry the same PTM will necessarily follow the same abundance pattern. Therefore, it can occur that a significant effect at the PTM-level stems only from one or a few associated peptidoforms while the other associated peptidoforms remain unchanged between conditions. This might indicate that the underlying biology is not only affected by a single PTM, but rather by a combination of PTMs and/or sequence variation. We thus recommend adding a peptidoform analysis by default to the overall workflow.

Peptidoform level information was available in both simulations, hence the performance of our method can be evaluated at this level as well. The peptidoform level tpr-fdp plots are given in figures 5 and 6, and the underlying data in supplementary table 3 and 4. These show that msqrob2PTM also performs well on the peptidoform level and maintains good control of false positives. However, on peptidoform level, the method performance seems to be more affected by a lower number of replicates, increased variability, and missingness. This can be expected as there is inherently less information, but more variation, present at the peptidoform level. This variability is reduced by averaging over peptidoforms when summarising the data to the PTM level. However, because PTMs are not directly quantified, but averaged out over peptidoforms, they can lead to more ambiguous results.

**Figure 5:**
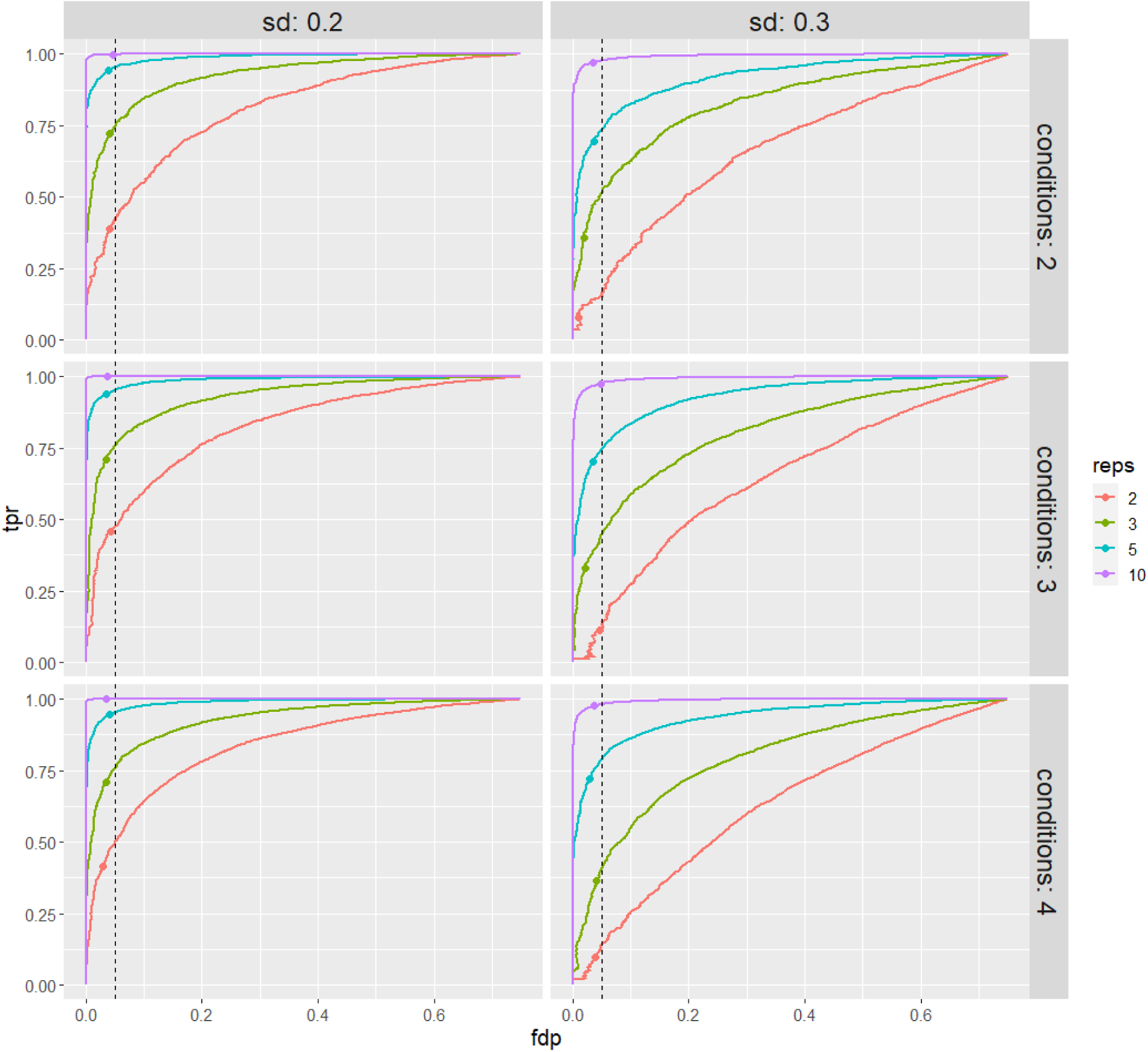
tpr-fdp plot for datasets simulated under the first scenario (no missingness). Performance of msqrob2PTM is assessed at peptidoform level. Observed fdp at a 5% FDR cut-off is denoted by dots.

**Figure 6:**
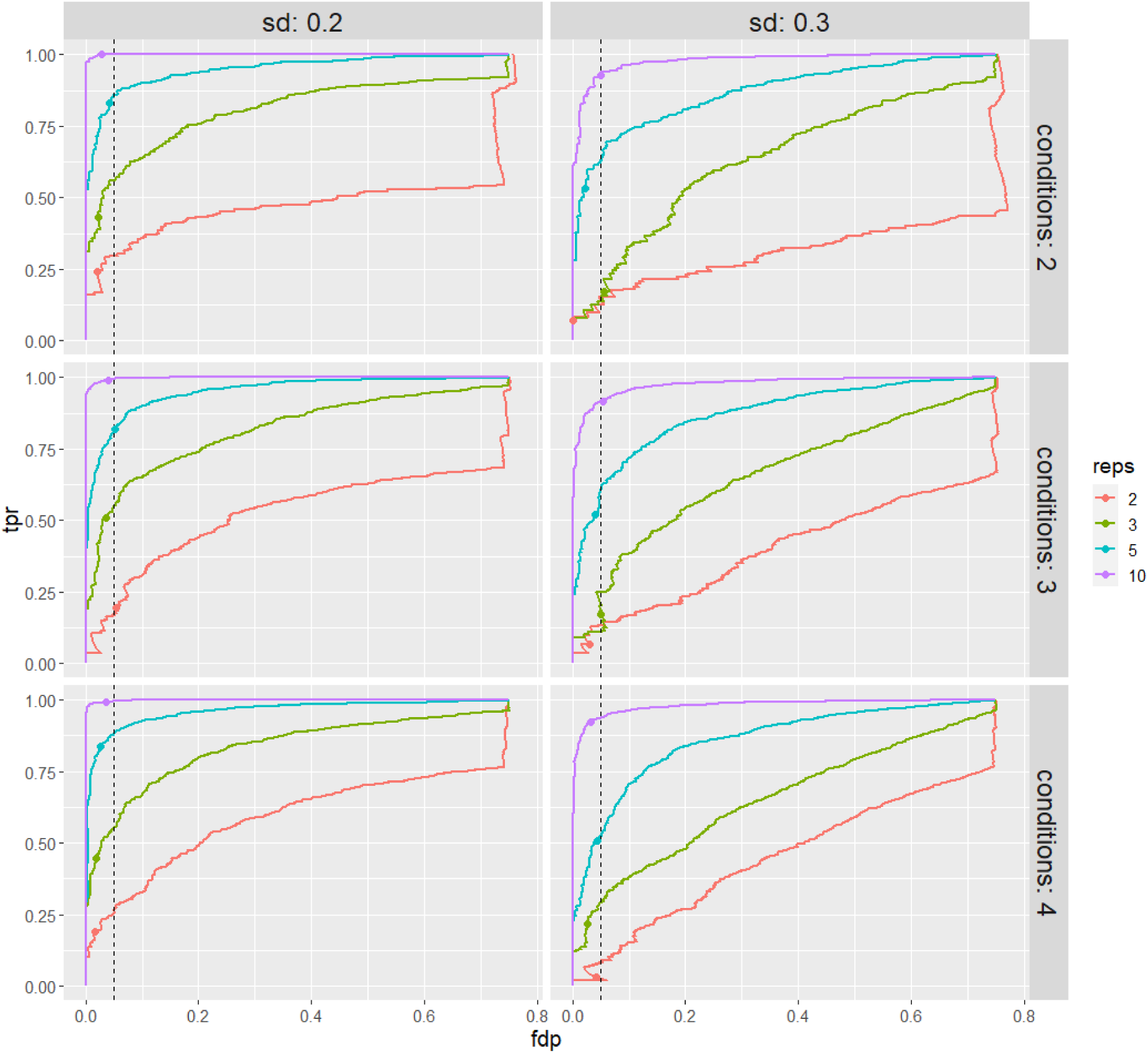
tpr-fdp plot for datasets simulated under the second scenario (with missingness). Performance of msqrob2PTM is assessed at peptidoform level. Observed fdp at a 5% FDR cut-off is denoted by dots. For datasets with only 2 or 3 replicates, the method starts to suffer from lack of information, making it harder to report significant peptidoforms, especially for datasets with sd 0.3.

Note that, as MSstatsPTM does not offer a peptidoform level analysis, no comparison could be included for this workflow.

### Biological spike-in dataset

The design of the spike-in dataset (see also figure 2) is suboptimal to assess the performance of methods inferring differential PTM usage. This is because the spiked-in peptides and their corresponding protein abundance in the background proteome are not correlated as they would be in real experiments. Indeed, the latter does not contain the actual parent proteins of the spike-in peptides. Moreover, the E.coli proteins in mixes 3 and 4 induce loading differences present across the samples (see also supplementary figure 3), which brings additional normalisation issues. We illustrate these issues using ROC curves that compare the performance of different approaches: differential PTM abundance by adopting a conventional msqrob2 workflow directly on the summarized PTM-level intensities without normalisation (DPA-NonNorm), the same workflow upon normalisation with the median peptidoform log-intensity (DPA), the default workflow for msqrob2PTM (default msqrob2PTM workflow assessing DPU), and MSstatsPTM (default MSstatsPTM workflow) (figure 7). Every pairwise comparison between mixes is shown. Because all methods report many false positives for this dataset, the tpr-fdp plots quickly became unreadable (see supplementary figure 4).

**Figure 7:**
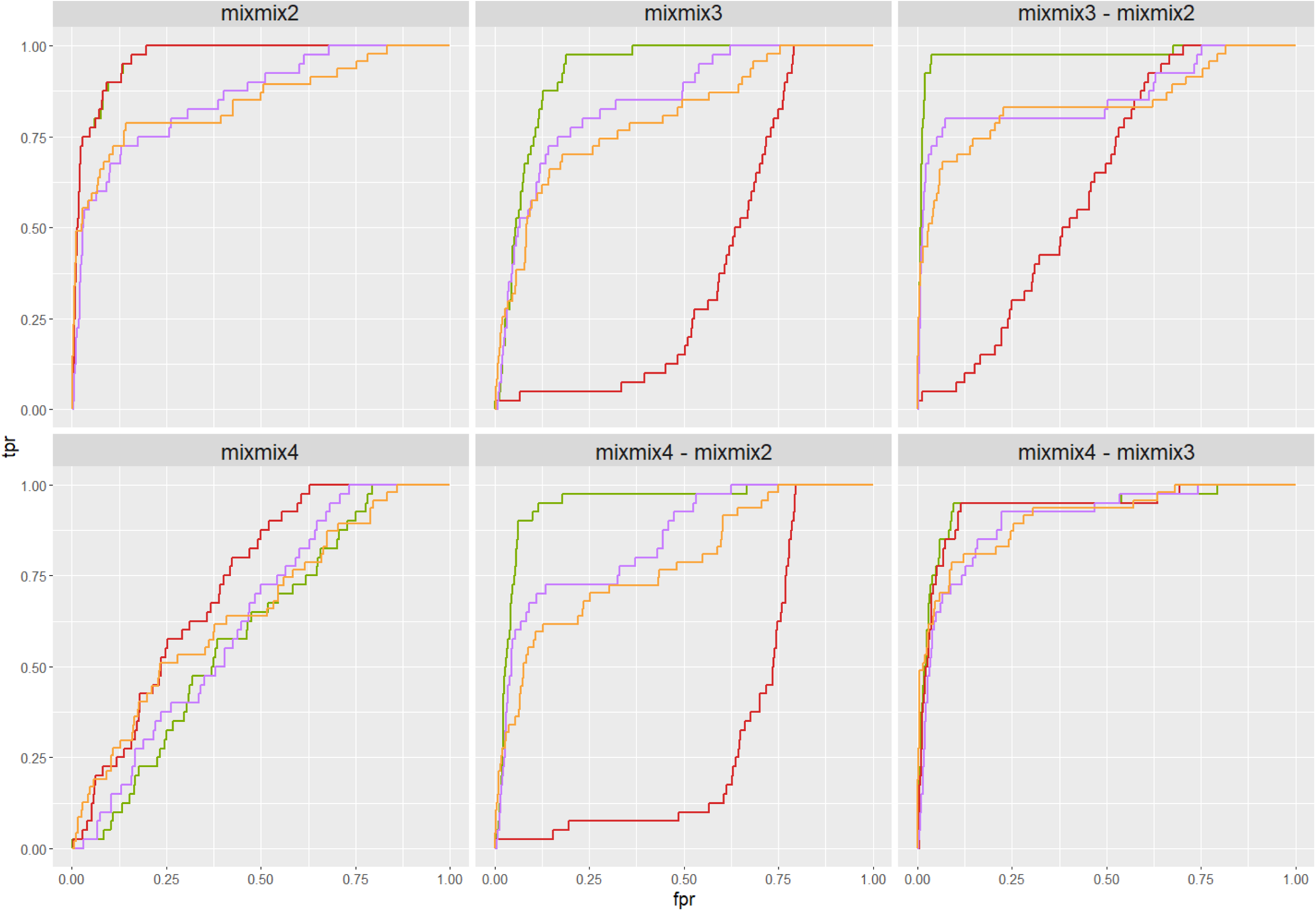
ROC curves of the different approaches for all pairwise comparisons. Mix 4 vs 1 (mixmix4) serves as internal control, thus the curves should follow the diagonal as closely as possible. DPA performs very well in all comparisons and outcompetes all other methods. DPA-NonNorm has good performance in the two comparisons where adjusted and unadjusted fold changes are the same, but breaks down for the other comparisons. The performance of MSstatsPTM and msqrob2PTM (the default differential PTM usage workflow) is similar, with performance dependent on the comparison being made.

When comparing mix 4 to mix 1 (mixmix4), the log2FC after adjustment should be 0, hence, no method should report any differential PTMs. Indeed, this comparison is an internal control, and the ROC curves are expected to lie along the diagonal. Here, *DPA-NonNorm* and *MSstatsPTM* show the largest deviations from the diagonal.

In the other comparisons, DPA always outperforms the other methods. Note that DPA assesses differential PTM abundance rather than differential usage as it does not normalise for parent protein intensity. This superior performance of the DPA method as compared to the DPA-NonNorm method indicates that it is very important to correct for technical variability resulting from the experimental design, i.e. the loading differences, rather than correcting for parent protein abundance. In the mix 2 vs mix 1 (mixmix2) and mix 4 vs mix 3 comparison, DPA-NonNorm also performs very well, because in these comparisons, the adjusted and unadjusted fold changes are the same. However, the loading differences for the other comparisons cause a breakdown of DPA-NonNorm. MSstatsPTM and msqrob2PTM always have a lower performance than DPA, but never break down. For the mix 2 vs mix 1 and the mix 4 vs mix 3 comparisons, MSstatsPTM performs slightly better than the default msqrob2PTM workflow, while the latter performs better in the remaining three comparisons. The decrease in performance by msqrob2PTM as compared to DPA can be explained by the increase in variability that is introduced in the workflow by subtracting the unrelated “parent protein intensities” from the spiked-in peptidoform intensities. In other words, the design is not suited to benchmark the performance of methods developed to quantify differential peptidoform usage. However, the design is useful for assessing the performance of methods that quantify differential PTM abundance. This can easily be obtained with standard msqrob2 workflows, but is not returned by default by MSstatsPTM. However, because the msqrob2 suite builds upon the QFeatures architecture, the results of a DPA and DPU workflow can both be stored in the same object, thus providing more transparency and reproducibility across the workflows.

For completeness, we also plotted the log2 fold changes for all PTMs in supplementary figures 5 and 6, which illustrate that both msqrob2PTM as well as MSstatsPTM provide good estimates for these.

### Ubiquitination dataset

msqrob2(PTM) is capable of handling more complex designs that require mixed model analysis, as well as datasets that lack a non-enriched version of the dataset. These two aspects apply to the ubiquitination dataset. Note that this is an experimental, biological dataset, and therefore does not come with a known ground truth.

Despite the two abovementioned complexities, the standard msqrob2PTM workflow could find differentially abundant ubiquitin sites in most comparisons, except for the USP30_OE vs control comparison. However, table 2 shows that msqrob2PTM generally reports much fewer significant PTMs than MSstatsPTM.

**Table 2:** The number of significant PTMs reported for each contrast for both methods.

| Contrast | MSstatsPTM | Msqrob2PTM |
| --- | --- | --- |
| Combo vs Ctrl | 424 | 30 |
| CCCP vs Ctrl | 359 | 12 |
| USP30_OE vs Ctrl | 40 | 0 |
| Combo vs CCCP | 31 | 1 |
| Combo vs USP30_OE | 407 | 24 |
| CCCP vs USP30_OE | 364 | 13 |

Upon closer inspection of the PTMs reported as significant by MSstatsPTM, it was discovered that this large discrepancy can be explained by several reasons.

First, both methods have a different way of dealing with missing data. Upon inspecting multiple line plots, we observed PTMs that were flagged as significant by MSstatsPTM despite having only one bio-repeat, or even only a single data point available in one of the conditions. In figure 8, for instance, line plots are shown for two PTMs that are significant in MSstatsPTM when comparing the combination condition (Combo) *versus* the control condition (Ctrl), but not in msqrob2PTM. Notably, PTM O00154_K205 only presents PTM information for the first biological replicate, while PTM O00159_K0578 contains just one data point within the entire control condition. For these features, msqrob2 therefore did not return a model fit.

**Figure 8:**
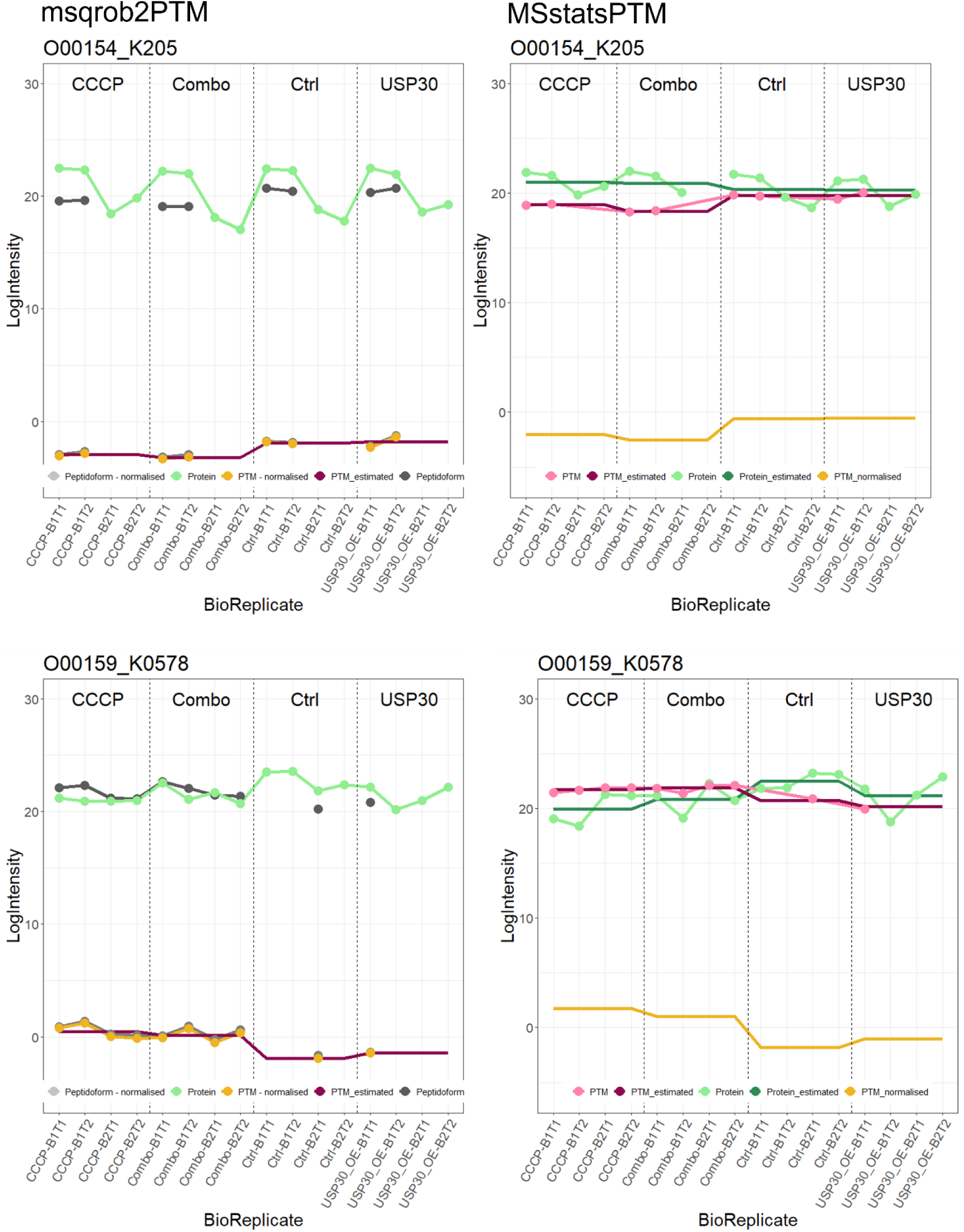
line plots displaying estimated log_2_ intensity values of the PTM (dark pink) for each sample, its normalised intensity values (yellow), log_2_ intensity values of its parent protein (green), for MSstatsPTM estimated log_2_ intensity values of that parent protein (dark green), and for msqrob2PTM, log_2_ intensity values of the peptidoforms (grey) on which the PTM occurs. On the left, line plots for PTM O00154_K205 and O00159_K0578 for msqrob2PTM, on the right for MSstatsPTM. Both PTMs were deemed significant by MSstatsPTM when comparing the control condition to the combination condition (combo), but not by msqrob2PTM. O00154_K205 only contains intensity information for bio replicate B1. O00159_K0578 only has 1 associated intensity value in the control condition. Hence, both of these PTMs contain too few datapoints for msqrob2PTM to determine significance.

When examining the results more closely, we noticed that MSstatsPTM uses three different models to fit the data and that the model choice is based on the available data points for each PTM (see UbiquitinationBioData exploration of results file on https://github.com/statOmics/msqrob2PTMpaper for detailed examples), i.e. a full mixed model was employed when no data was missing, using a fixed effect for group and random effects for subject (1 | SUBJECT) and subject x group (1 | GROUP:SUBJECT), as soon as a single data point is missing, the (1 | GROUP:SUBJECT) term is dropped, and when data is missing for one of the bio repeats in all conditions, a linear model is employed with only a fixed group effect. This adaptability to missing data comes with a price, however. Notably, the second model, without the (1 | GROUP:SUBJECT) term, ignores the between bio repeat variability. Indeed, bio repeat 1 in the control group is not the same as bio repeat 1 in the combination group. However, they are treated as such, resulting in underestimated standard errors.

Across comparisons, 15-27% of PTMs deemed significant were modelled with an incorrect mixed model (% differs according to comparison). Moreover, 44-75% of significant PTMs were modelled using a linear model, which represents features for which msqrob2 does not fit any model at all because biological repeats are lacking. Moreover, when examining the significant PTMs together with their parent proteins, it became apparent that for most features the PTM and protein intensities were modelled with a different model. This can lead to artifacts such as shown in figure 8 (top panels), where the protein data contains information about only one of the two bio repeats, but is still used to make the adjustment for the other bio repeat! To avoid these ambiguities, we conducted an MSstatsPTM-like analysis while enforcing the use of the full mixed model. Only PTMs with associated parent proteins were included in the analysis. Subsequently, the full mixed model was applied to both the PTM and protein-level data. The adjustment for protein abundance followed the standard MSstatsPTM procedure, and the resulting p-values were adjusted using the Benjamini-Hochberg method. Using the native MSstatsPTM implementation the “CCCP” vs “Ctrl” comparison identified 359 significant PTMs. However, when solely employing the full mixed model, only 55 PTMs remained significant, which is in line with our msqrob2 results.

Second, the two methods employ distinct conceptual approaches. In msqrob2PTM, within-sample normalisation according to protein level abundance is performed first, followed by statistical analysis. MSstatsPTM, however, uses the modelled PTM and protein results for normalisation, ignoring the inherent biological correlation between PTMs and their parent proteins within a sample. Analysing these separately can sometimes generate ambiguities. Figure 9 illustrates this issue, demonstrating a PTM that was flagged as significant for the “Combo” vs "Ctrl” comparison by MSstatsPTM, but not by msqrob2PTM. Specifically, the peptidoform carrying PTM O60260_K369 closely mirrors the intensity pattern of its parent protein, resulting in minimal differences, and therefore no significant regulation, in PTM intensities after normalisation for protein abundance in our msqrob2PTM workflow. However, as MSstastPTM first fits models to the PTM and protein level data separately, and only afterwards uses these model estimates to correct for the difference in protein abundance, differences in PTM usage are artificially enlarged, leading to a significant PTM according to MSstatsPTM in this comparison.

**Figure 9:**
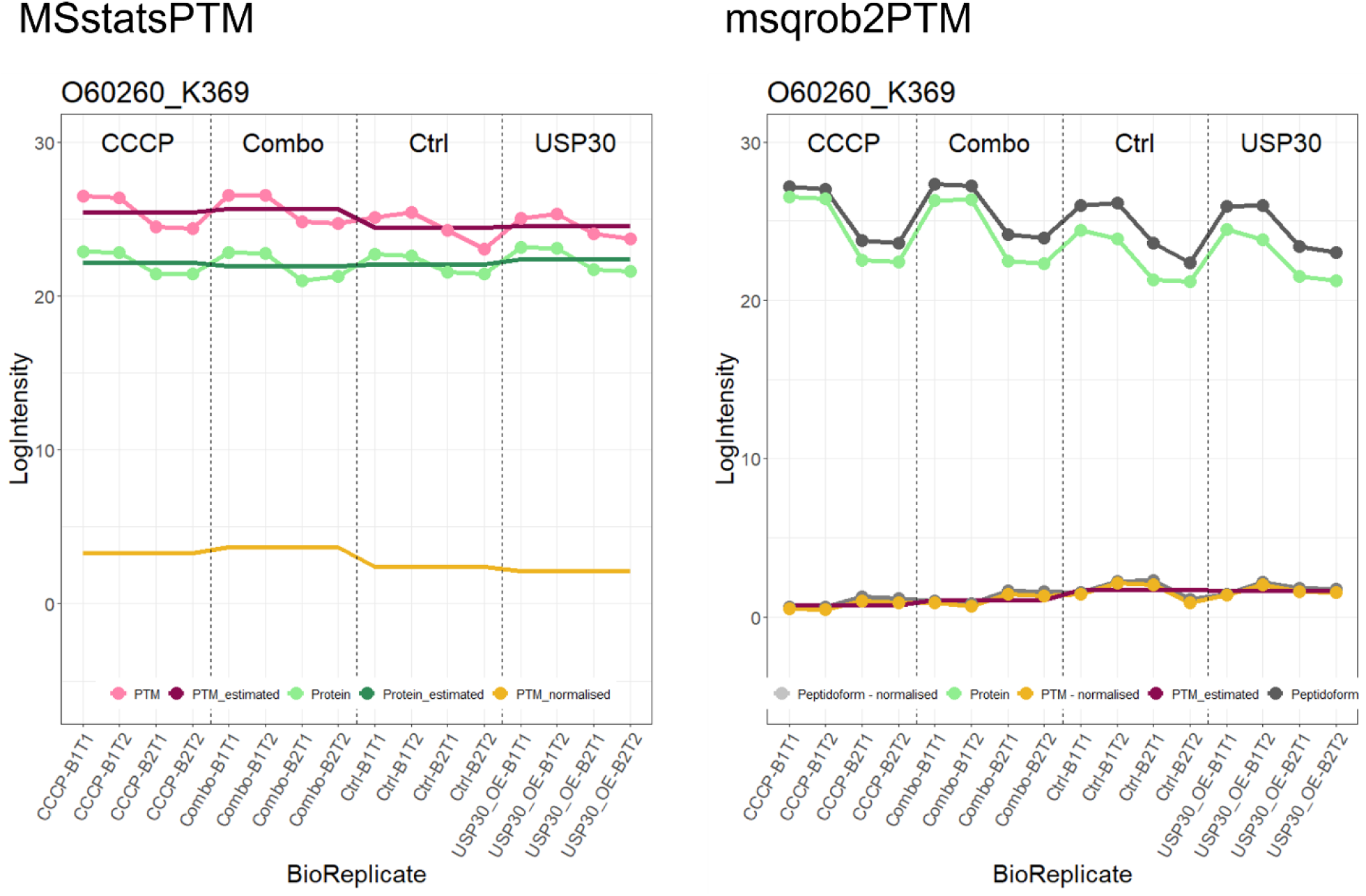
line plot displaying PTM log_2_ intensity values (pink dotted line) or peptidoform log2 intensity values (dark grey dotted line) and log_2_ intensity values of its parent protein (light green dotted line) in each sample. MSstatsPTM first fits a model to PTM (dark pink line) and to protein intensities (dark green line) to estimate average intensity in each condition. Subsequently, fitted average protein abundances are subtracted from fitted average PTM intensities to obtain average PTM abundances in each condition corrected for protein abundance (yellow line). Conversely, msqrob2PTM first normalises peptidoform intensities using parent protein abundance, resulting in a normalised peptidoform (light grey dotted line). From normalised peptidoforms, normalised PTM intensities are calculated (yellow dotted line). Estimated log2 intensity values of the PTM are depicted in dark pink. MSstatsPTM corrected PTM abundances seem to indicate differential PTM usage. Moreover, the comparison between “Combo” vs “Ctrl” is returned by MSstatsPTM as statistically significant. This, however, appears to be an artifact of MSstatsPTM as the correction for protein abundance does not account for the link between protein and PTM intensities within-sample. Indeed, when comparing “Combo” and “Ctrl” sample level intensities, the pattern at PTM-level closely follows that of its parent protein.

### Phospho dataset

Two different workflows were employed for this dataset. The first workflow uses the non-enriched counterpart dataset to normalise for differences in protein abundance, while the second workflow only used the enriched dataset, also for the normalisation step. It is important to note that two distinct instrument platforms were used to analyse the total proteome and phosphoproteome samples. The chromatographic conditions were identical as well as the MS instrument geometry but two consecutive generations of Q-Orbitraps were used (Q-Exactive Plus versus Q-Exactive HF-X). This partly explains the observed heterogeneity between enriched and non-enriched datasets. Indeed, we observed a substantial proportion (approximately 25%) of proteins present in the enriched dataset that were absent in the non-enriched one. This led to some PTMs that could not be normalised, which we opted to exclude from subsequent analysis in workflow 1.

Both workflows involved testing multiple contrasts based on two factors: condition (A or B), and subset (x or y). In the first workflow (utilising both datasets), 31 unique differential PTMs were found, of which 25 phosphorylations. Most of these PTMs exhibited significant downregulation in condition A compared to B within subset y.

In the second workflow (using only the enriched dataset), fourteen unique significant PTMs were identified, of which eight phosphorylations. The majority of phosphorylations showed significant differential usage between condition A and B within subset y and/or exhibited significant differential usage between condition A and B averaged over subsets x and y. Supplementary tables S5 and S6 provide detailed results.

Interestingly, the results differ between the two workflows. Of the 31 PTMs identified in workflow 1, ten were also found in workflow 2.

Instead of solely focusing on significant PTMs, our method is capable of detecting differentially used peptidoforms as well. For this dataset, the first workflow detected twelve peptidoforms as differentially abundant, predominantly showing downregulation in condition A for subset y.

In the second workflow, which lacked a global profiling dataset, seven significant peptidoforms were detected across the different comparisons. LPIVNFDYS[Phospho (STY)]M[Oxidation (M)]EEK was picked up as DU by both workflows and is particularly interesting, because both PTMs present on this peptidoform are also returned as significant in the differential PTM usage analysis. Hence, one of the PTMs might have been detected as differential because the other PTM is also present on the same peptidoform, potentially influencing its significance upon averaging with the remaining peptidoforms carrying this PTM. To assess the contribution of different peptidoforms to a single PTM, line plots can be used to visualise both the PTM intensities across the samples as well as the intensities of its contributing peptidoforms. Figure 10 illustrates this issue. Indeed, the top panel shows a phosphorylation that occurs in two peptidoforms, the bottom panel shows an oxidation that also occurs on one of these peptidoforms. The peptidoform with both modifications was significant, while the second peptidoform that did not carry the oxidation was not significantly DU. The intensity for the phosho-PTM is obtained upon summarisation over both peptidoforms, and was reported significant when assessing the data at the PTM-level. However, the significance of the phospho-PTM might be an artifact triggered by the presence of additional oxidation in one of its underlying peptidoforms.

**Figure 10:**
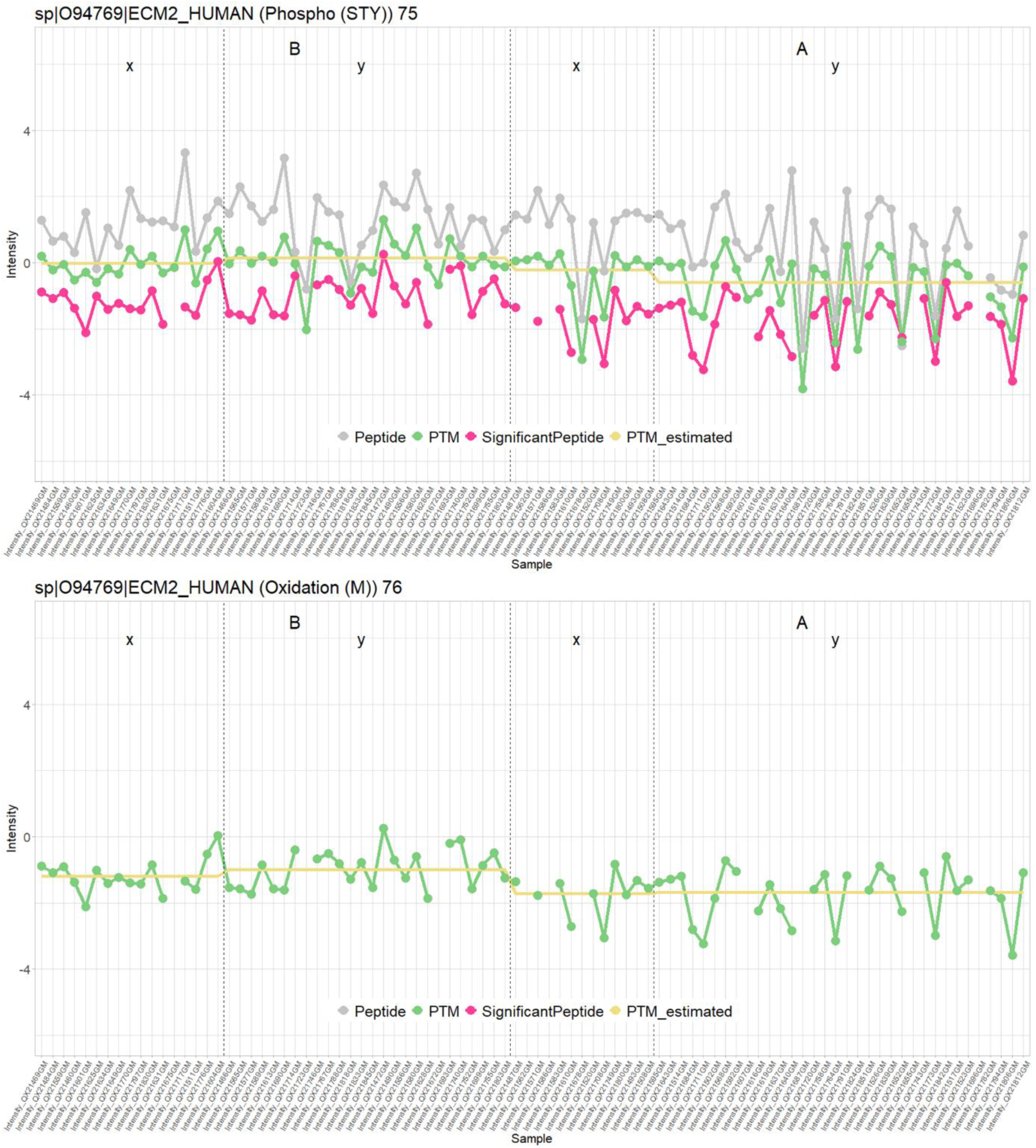
Line plots of normalised intensity values per sample for significant peptidoform (LPIVNFDYS[Phospho (STY)]M[Oxidation (M)]EEK) and its corresponding PTMs for the phospho dataset. At the top, the significant peptidoform is depicted in pink. In green is the PTM occurring on that peptidoform, in this case phosphorylation. In grey any other peptidoform carrying that same PTM, and in yellow, the PTM intensity value as estimated by the model. The PTM is represented by two peptidoforms that roughly follow the same pattern, resulting in a PTM that resides in the middle. At the bottom we see the other PTM occurring on that peptidoform, the oxidation. No other peptidoform carries that same modification, resulting in perfect overlap between the line of the significant peptide and that of the PTM. Here, it is possible that the oxidation is only significant because the phosphorylation is. Indeed, the driving force of the significance of this particular peptidoform could be coming from the phosphorylation (which has two associated peptidoforms). Note that, while these particular line plots were derived using the workflow without a non-enriched dataset, the corresponding plots from workflow 1 are extremely similar.

Some PTMs are also significant because they enable aggregating evidence over multiple non-significant peptidoforms that all have a similar expression pattern. An example of this can be seen in figure 11 for sp|P10451|OSTP_HUMAN (Phospho (STY)) 280.

**Figure 11:**
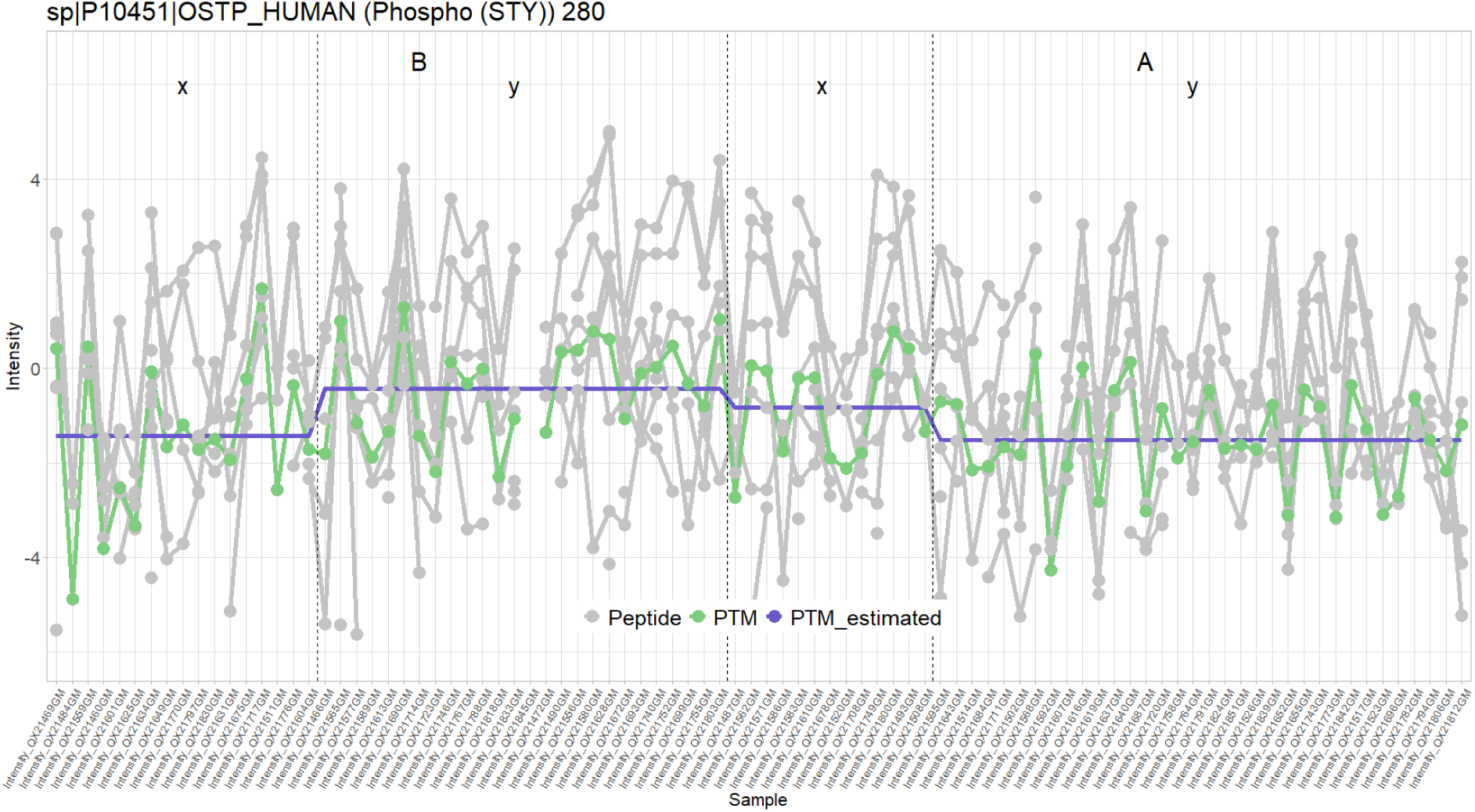
Line plot of normalised intensity values of significant PTM sp|P10451|OSTP_HUMAN (Phospho (STY)) 280 and its associated peptidoforms. In green the summarised and normalised intensity value of the PTM, in grey all peptidoforms (normalised) containing this PTM, in purple the PTM intensity values as estimated by the model. While none of the peptidoforms are individually significant, these all contribute to a PTM that can be picked up as differentially abundant (downregulated in condition A for samples from subset y).

### Mock analyses

As the phospho datasets are biological experiments, the ground truth is unknown. Therefore, we cannot assess the performance of each method. We also do not know if the method provides reliable false positive control. To assess if our workflows provide good type I error control for the case study, we therefore perform a mock analysis. In particular, we introduce a factor for a non-existing effect, implying that all features that are returned significant upon testing for this factor are false positives. Here, we focus on subset y from condition B, so that ample samples remain. When the method provides good false positive control, the p-values upon assessing the mock effect will be uniform.

The p-value distribution for the workflow that only uses the enriched dataset is given in Figure 12. The top panels show the results for the PTM-level analysis and the bottom panels for peptidoform analysis. Both workflows with and without robust regression provide fairly uniform p-values. Supplementary figures 7-10 show similar plots for four other random mock datasets, showing consistency of performance.

**Figure 12:**
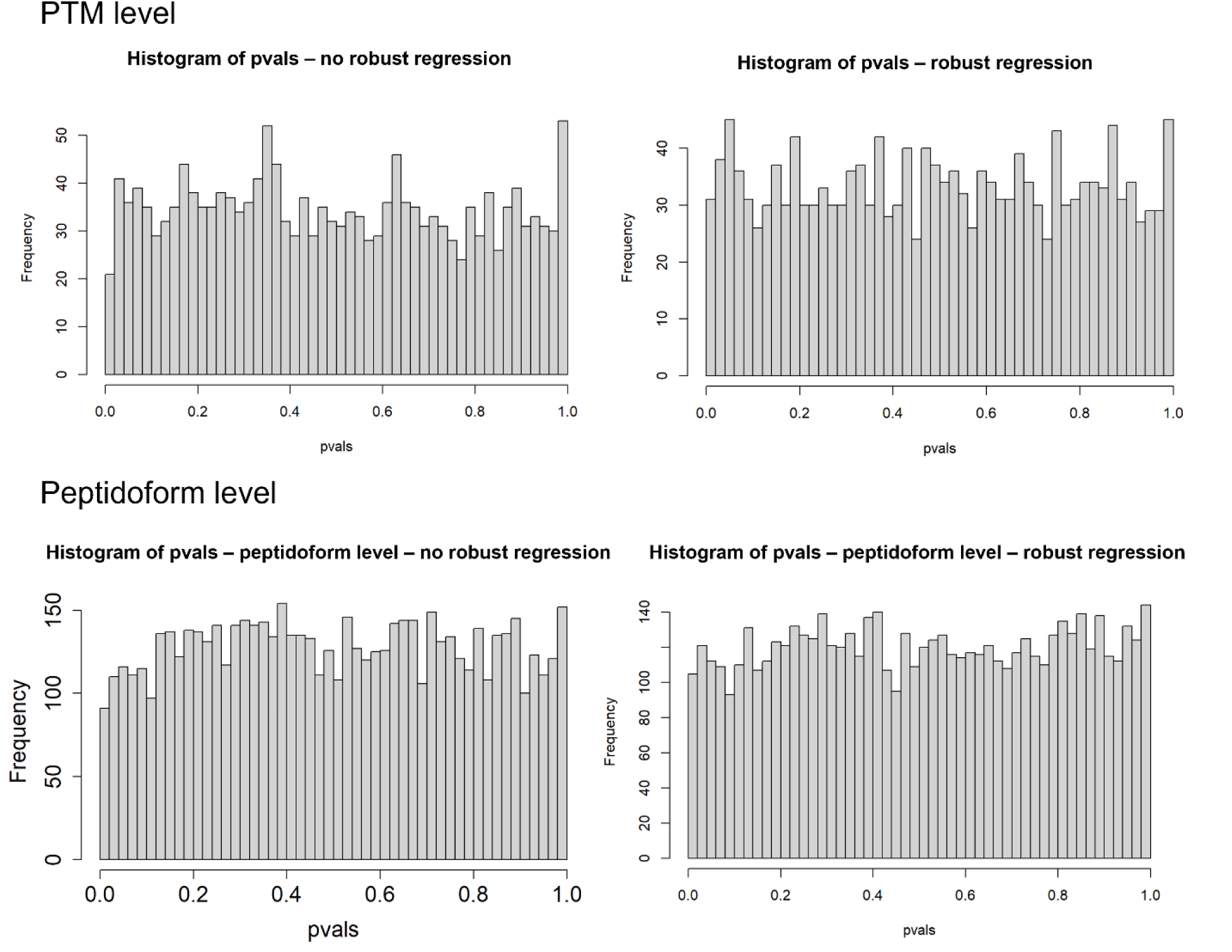
Distribution of p-values for mock analysis of the phospho dataset without global profiling run, for analysis on PTM level (top) as well as peptidoform level (bottom). Left panels are for workflows without robust regression in the modelling step; Right panels correspond to workflows with robust regression in the modelling step. All p-values are fairly uniform, indicating acceptable type I error control.

We did a similar mock analysis for the workflow that uses the non-enriched dataset for usage calculation (Figure 13). The workflow on peptidoform level using robust regression showed a slight increase in low p-values, which is also observed in some other random mock datasets (Supplementary Figures 11-14). The remaining workflows generated fairly uniform p-values for all random mock datasets (Figure 13 and Supplementary Figures 11-14). We therefore did not adopt robust regression for the peptidoform analysis.

**Figure 13:**
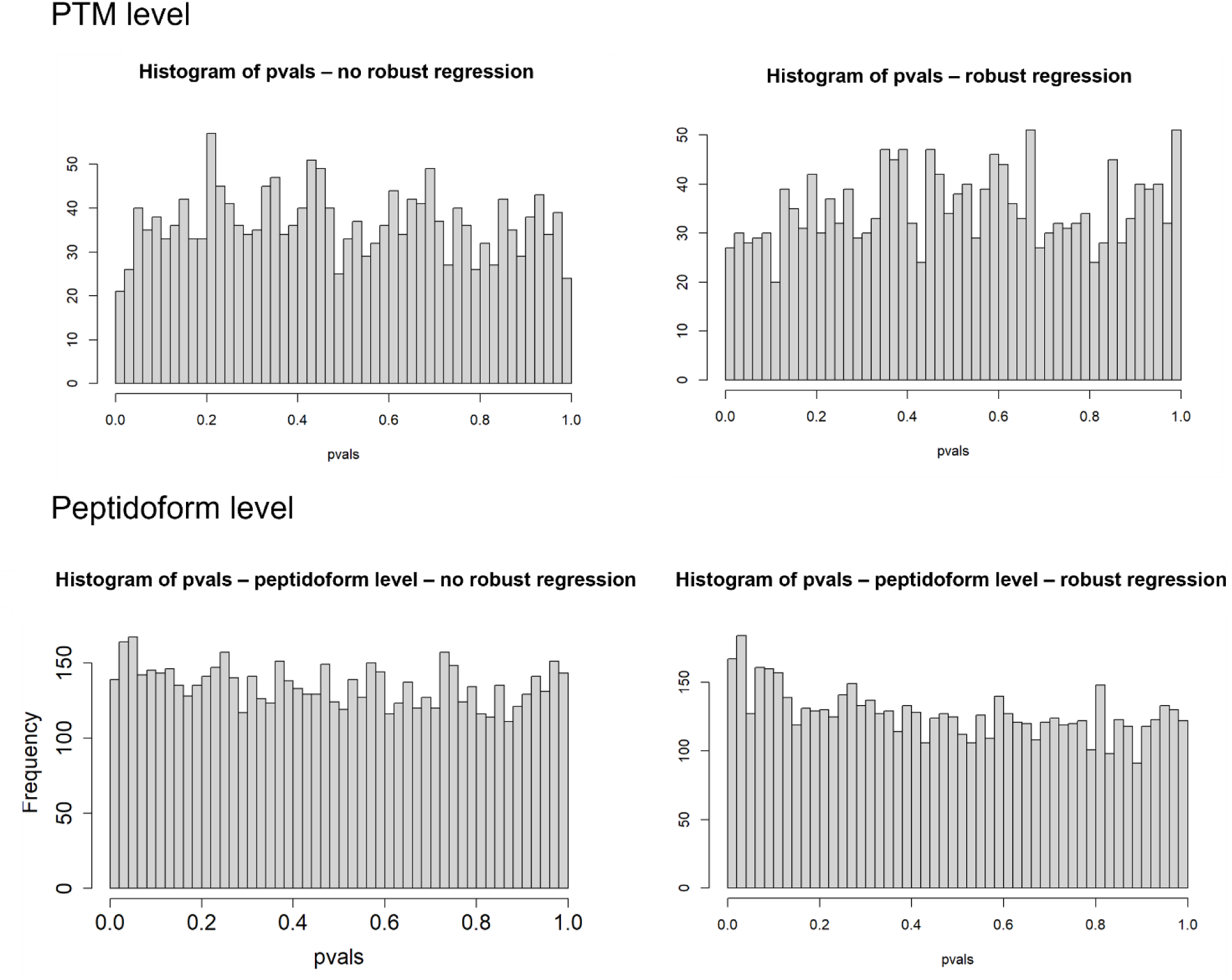
Distribution of p-values for mock analysis of the phospho dataset using the non-enriched dataset to estimate the usages. Results at PTM level (top panels) as well as at peptidoform level (bottom panels). Left panels are based on a workflow without robust regression; right panels on a workflow with robust regression.

## Discussion

We here introduced msqrob2PTM, a novel workflow in the msqrob2 universe, designed for performing differential abundance as well as usage analysis on PTM and peptidoform level. These two analyses are distinguished by their normalisation strategies. In abundance analysis, only a normalisation to reduce technical variation is included, while the novel usage workflow incorporates normalisation against parent protein intensities. Both approaches have their relevance in PTM research. DPU enables the discovery of differential PTMs that respond differently than their parent protein. However, in certain scenarios, DPA might be of interest instead. Indeed, when an increase in total protein concentration leads to a corresponding increase in PTM concentration, there may be biological implications associated with this elevation in PTMs, regardless of whether it is driven by changes in parent protein levels or not. Therefore, the choice between DPA and DPU depends on the specific research question at hand, or they can both be performed to complement each other.

Through analysis of simulated and biological datasets, we have demonstrated that our workflows improve upon the state-of-the-art MSstatsPTM. We showed the advantage of first normalising the peptidoform intensities by the parent protein abundance before conducting the differential analysis. In this way, we can immediately model the usages as opposed to MSstatsPTM that estimates the fold changes for the PTM and protein values separately before differencing these to estimate DPU. Indeed, the peptidoform and protein values from the same sample are correlated, which is explicitly accounted for in our DPU workflow but is ignored by MSstatsPTM. We showed for the latter method that this can lead to artifacts in the estimated fold change for some PTMs upon correction for the fold change in the parent protein. Moreover, MSstatsPTM also ignores the correlation when calculating the variance on the difference in fold change leading to incorrect inference.

Another key distinction between both packages is how they handle PTMs that cannot be fitted with the desired model. MSstatsPTM prioritises automation and aims to infer on as many PTMs as possible. However, this leads to reporting on PTMs for which the fit is based on different models and often on insufficient data to draw reliable inference on the contrast of interest. Moreover, for PTMs that lack a corresponding protein expression fold change, results are returned based on the PTM fold change alone. Hence, MSstatsPTM silently combines inference on differential usage with inference on differential abundance in one output list depending on the degree of missingness at the protein-level. In general, a standard user is not fully aware of these issues, and the subtleties of interpretation that these require. In contrast, our msqrob2PTM workflow emphasises transparency and reproducibility. While this choice may lead to some PTMs that cannot be estimated using the default workflow, it does ensure that users are fully aware of what was modelled for each PTM. Moreover, we feel that PTMs for which no results are returned due to missingness require the intervention of a skilled data analyst to develop tailored solutions to infer on differential abundance and/or usage; solutions that are moreover supported by the msqrob2 universe. Indeed, we showed that automatic approaches can lead to biased results, and especially in experiments with more complex designs.

These differences in normalisation approach and design concept elucidate the variations in performance across the different datasets that were used in our benchmark. In the simulated datasets, msqrob2PTM capitalises on the within-sample correlation between peptidoforms and proteins that is present in the data, resulting in superior performance compared to MSstatsPTM. However, in the spike-in dataset, where this correlation is absent due to its unrealistic design, the default msqrob2PTM workflow exhibits similar performance to MSstatsPTM. However, for this dataset we show that our workflow for assessing differential PTM abundance analysis uniformly outperforms both the msqrob2PTM and MSstatsPTM workflows assessing differential PTM usage. Indeed, the spike-in study is suited for assessing the performance on differential PTM abundance rather than on differential PTM usage, as the spiked PTMs were not correlated to their corresponding protein in the background. In the biological ubiquitination dataset, the high amount of missing data, and the absence of a global profiling dataset leads to a high number of PTMs that cannot be fitted with the required model. MSstatsPTM will then resort to other, simpler models that are often suboptimal or even mismatched, while msqrob2PTM will simply not return results for these PTMs, leading to a lower number of reported significant PTMs.

These datasets bring to attention a broader issue in the field, specifically the scarcity of suitable datasets for accurately assessing Differential Peptidoform Usage (DPU). When designing such experiments, it is favourable to incorporate a global profiling dataset along with an adequate number of biological replicates. This comprehensive approach not only enables a more thorough evaluation of DPU but also enhances statistical power, yielding more reliable and robust results. Indeed, the approach benefits from multiple replicates per feature. As PTMs usually appear low abundantly, this is often challenging to achieve in practice (26).

Although we recommend the addition of a global profiling counterpart to an enriched PTM dataset, this is conceptually not required as normalisation can be done using all peptidoforms mapping to the same protein. However, we showed that this approach has the risk of partially diluting the effect of the PTM as their underlying peptidoforms are now involved in the calculation of the PTM usage.

As opposed to MSstatsPTM we do not make use of converters. Hence, msqrob2 input is not restricted to certain search engines or quantification algorithms, providing the user with full flexibility. However, this does require the user to convert their data into appropriate input format, which is a simple flat text file format (as exportable from a spreadsheet) or a data frame in R that can be used by the constructor for QFeatures objects. Furthermore, our workflows are modular and provide the user with the flexibility to use custom pre-processing steps. Default workflows are presented in our package vignettes, but these can easily be altered by building upon methods in the QFeatures package. Moreover, the use of the QFeatures infrastructure also guarantees that input data is never lost during processing, but remains linked to the pre-processed and normalised assays as well as to the model output, insuring transparency, traceability, and reproducibility. This allows the user to perform differential usage (and/or abundance) analysis on both PTM and peptidoform (or even protein) level, while storing and linking all these different results in a structured manner in the same object.

Another advantage of msqrob2PTM is that it can manage multiple modification sites per peptidoform. The peptidoform will then simply be used in the summarisation of multiple PTMs. This is particularly useful when using open modification search engines, which can often find multiple PTMs per peptide. Moreover, we also include workflows on differential abundance and usage analysis on the peptidoform level. Indeed, as shown in figures 10 and 11, it can be relevant to know whether a significant PTM stems from multiple (slightly) significant associated peptidoforms, or whether it is driven by one or a few very strongly significant associated peptidoform(s). In the latter case, it could be possible that these significant peptidoforms carry another modification that is driving the differential usage. Hence, we always advise users to conduct a peptidoform level analysis as well.

Overall, we have shown that our msqrob2PTM workflow is a sensitive and robust approach compared to the state-of-the-art, while providing good fpr control and high accuracy. Our modular implementation offers our users full flexibility with respect to the search engine and pre-processing steps, while still offering a comprehensive, transparent, and reproducible workflow that covers the entire differential PTM analysis.

## Code and data availability

The analysis files and data are available on https://github.com/statOmics/msqrob2PTMpaper and PRIDE PXD043476.

## Supplementary materials

This article contains supplemental data.

## Acknowledgements

This research was funded by the Research Foundation Flanders (FWO) as a mandate awarded to **ND** (1S77220N), and as project funding awarded to **LM** (G010023N, G028821N) and **LC** (G062219N), funding from the European Union’s Horizon 2020 Programme to **LM** (H2020-INFRAIA-2018-1) [823839], and funding from a Ghent University Concerted Research Action to **LM** [BOF21/GOA/033] and **LC** [BOF20/GOA/023]. **LCG** and **PL** were supported by the Bundesministerium für Bildung und Forschung (01GM1917A, “Multi-omic analysis of axono-synaptic degeneration in motoneuron disease (MAXOMOD)”, which was funded in the scope of the E-Rare Joint Transnational Call for Proposals 2018 “Transnational research projects on hypothesis-driven use of multi-omic integrated approaches for discovery of disease causes and/or functional validation in the context of rare diseases.” Phosphoproteomics experiments were supported by the french proteomics infrastructure (ProFI FR2048, ANR-10-INBS-08-03).

## Author contributions

N.D.: writing, method development, data analysis

M.G.: writing, phosphorylation experiment and analysis

L.C.G.: phosphorylation experiment

P.L.: supervision phosphorylation experiment

C.C.: supervision phosphorylation experiment

L.C.: supervision, writing, method development

L.M.: supervision, writing

## Abbreviations

DPA: differential PTM abundance
DPU: differential PTM usage
fdp: false discovery proportion
FDR: false discovery rate
fpr: false positive rate
LC: liquid chromatography
MS: mass spectrometry
PTM: post-translational modification
tpr: true positive rate
ROC: receiver operating characteristic

