## Supplemental material for "msqrob2PTM: differential abundance and differential usage analysis of MS-based proteomics data at the post-translational modification and peptidoform level"

### Title

#### List of contents

#### Performance plots for the simulation studies

##### ROC curves simulation 1 for msqrobPTM and MSstatsPTM results

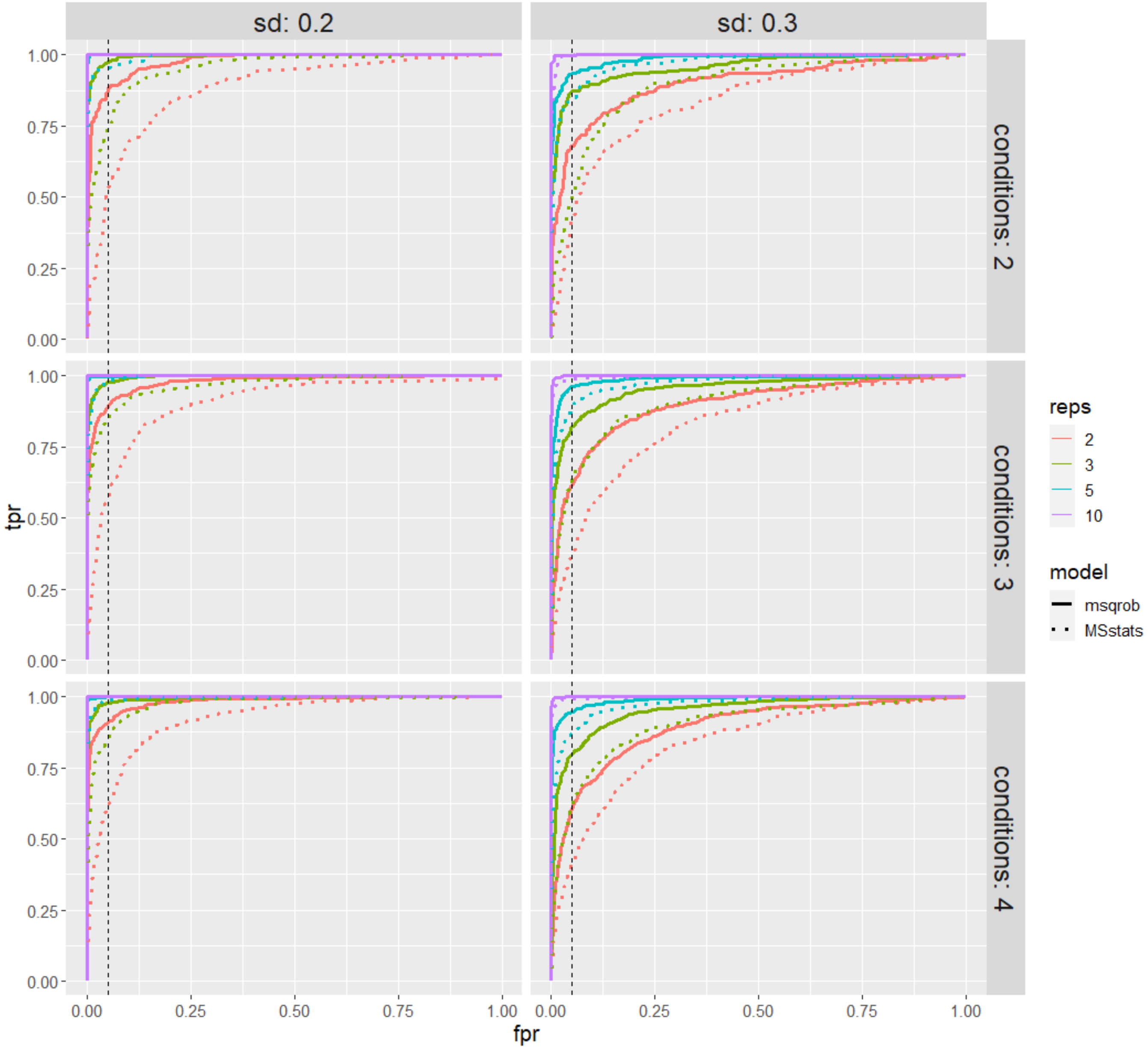

**Supplementary Figure 1:** ROC curves for the datasets of the first simulation. In solid lines are the ROC curves for msqrob, in dotted lines the ones for MSstats. The lay-out of the figure is the same as in supplementary figure 1, the difference is that on the x-axis the false positive rate is now plotted. Again, it can be observed that msqrob outperforms MSstats on all occasions.

#### ROC curves simulation 2 for msqrobPTM and MSstatsPTM results

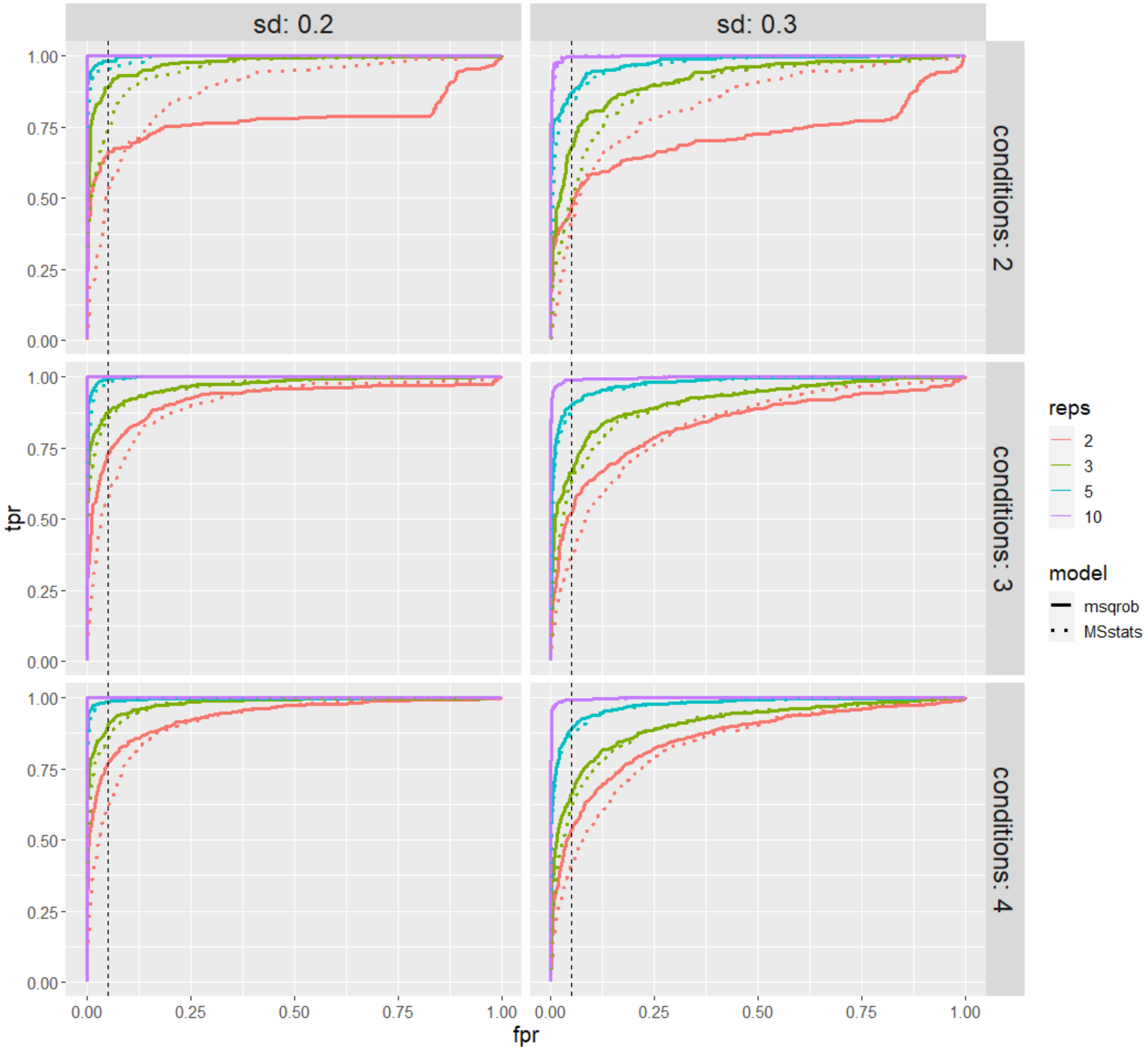

**Supplementary figure 2:** ROC curves for the datasets of the second simulation. In solid lines are the ROC curves for msqrob, in dotted lines the ones for MSstats. For the datasets with 2 conditions present, the MSstats curve ends above the msqrob curve. However, when looking at the 5% line (black dashed vertical line) in those plots, msqrob is always above its MSstats counterpart, indicating better performance when applying a 5% FDR cut-off.

### Performance metrics simulations msqrobPTM

#### Simulation 1 – PTM level

| SIMULATION 1 | | | | | | | | | |
|---|---|---|---|---|---|---|---|---|---|
| Dataset | sd | reps | conditions | fdp | sensitivity | specificity | precision | accuracy | fpr |
| 1 | 0,2 | 2 | 2 | 0,040 | 0,676 | 0,991 | 0,960 | 0,912 | 0,009 |
| 2 | 0,3 | 2 | 2 | 0 | 0,116 | 1 | 1 | 0,779 | 0 |
| 3 | 0,2 | 3 | 2 | 0,046 | 0,912 | 0,985 | 0,954 | 0,967 | 0,015 |
| 4 | 0,3 | 3 | 2 | 0,007 | 0,54 | 0,999 | 0,993 | 0,884 | 0,001 |
| 5 | 0,2 | 5 | 2 | 0,031 | 1 | 0,989 | 0,969 | 0,992 | 0,011 |
| 6 | 0,3 | 5 | 2 | 0,029 | 0,816 | 0,992 | 0,971 | 0,948 | 0,008 |
| 7 | 0,2 | 10 | 2 | 0,027 | 1 | 0,991 | 0,973 | 0,993 | 0,009 |
| 8 | 0,3 | 10 | 2 | 0,035 | 0,996 | 0,988 | 0,965 | 0,99 | 0,012 |
| 9 | 0,2 | 2 | 3 | 0,060 | 0,786 | 0,983 | 0,940 | 0,934 | 0,016 |
| 10 | 0,3 | 2 | 3 | 0,068 | 0,218 | 0,995 | 0,932 | 0,8005 | 0,005 |
| 11 | 0,2 | 3 | 3 | 0,053 | 0,93 | 0,983 | 0,947 | 0,9695 | 0,017 |
| 12 | 0,3 | 3 | 3 | 0,018 | 0,534 | 0,997 | 0,982 | 0,881 | 0,003 |
| 13 | 0,2 | 5 | 3 | 0,022 | 0,998 | 0,993 | 0,978 | 0,994 | 0,007 |
| 14 | 0,3 | 5 | 3 | 0,041 | 0,834 | 0,988 | 0,959 | 0,950 | 0,012 |
| 15 | 0,2 | 10 | 3 | 0,029 | 1 | 0,99 | 0,971 | 0,993 | 0,01 |
| 16 | 0,3 | 10 | 3 | 0,052 | 0,992 | 0,982 | 0,948 | 0,985 | 0,018 |
| 17 | 0,2 | 2 | 4 | 0,022 | 0,777 | 0,994 | 0,978 | 0,94 | 0,006 |
| 18 | 0,3 | 2 | 4 | 0,050 | 0,179 | 0,997 | 0,950 | 0,792 | 0,003 |
| 19 | 0,2 | 3 | 4 | 0,047 | 0,929 | 0,985 | 0,953 | 0,971 | 0,015 |
| 20 | 0,3 | 3 | 4 | 0,046 | 0,550 | 0,991 | 0,954 | 0,881 | 0,009 |
| 21 | 0,2 | 5 | 4 | 0,041 | 0,995 | 0,986 | 0,959 | 0,988 | 0,014 |
| 22 | 0,3 | 5 | 4 | 0,021 | 0,861 | 0,994 | 0,979 | 0,961 | 0,006 |
| 23 | 0,2 | 10 | 4 | 0,027 | 1 | 0,991 | 0,973 | 0,993 | 0,009 |
| 24 | 0,3 | 10 | 4 | 0,035 | 0,997 | 0,988 | 0,965 | 0,990 | 0,012 |

**Table S1:** Performance metrics for the PTM analysis of the 24 datasets in simulation 1. Essentially, the same conclusions can be drawn as from supplementary figure 1. MsqrobPTM generally performs very well, but performance can drop when the number of replicates get too low, especially when combined with a higher variation present in the dataset.

#### Simulation 2 – PTM level

| SIMULATION 2 | | | | | | | | | |
|---|---|---|---|---|---|---|---|---|---|
| Dataset | sd | reps | conditions | fdp | sensitivity | specificity | precision | accuracy | fpr |
| 1 | 0,2 | 2 | 2 | 0,024 | 0,346 | 0,820 | 0,976 | 0,702 | 0,003 |
| 2 | 0,3 | 2 | 2 | 0,024 | 0,176 | 0,830 | 0,977 | 0,669 | 0,001 |
| 3 | 0,2 | 3 | 2 | 0,034 | 0,676 | 0,990 | 0,966 | 0,911 | 0,008 |
| 4 | 0,3 | 3 | 2 | 0,071 | 0,368 | 0,987 | 0,929 | 0,832 | 0,009 |
| 5 | 0,2 | 5 | 2 | 0,048 | 0,956 | 0,984 | 0,952 | 0,977 | 0,016 |
| 6 | 0,3 | 5 | 2 | 0,016 | 0,728 | 0,996 | 0,983 | 0,929 | 0,004 |
| 7 | 0,2 | 10 | 2 | 0,038 | 1 | 0,987 | 0,962 | 0,99 | 0,013 |
| 8 | 0,3 | 10 | 2 | 0,050 | 0,98 | 0,983 | 0,950 | 0,982 | 0,017 |
| 9 | 0,2 | 2 | 3 | 0,057 | 0,432 | 0,967 | 0,943 | 0,834 | 0,009 |
| 10 | 0,3 | 2 | 3 | 0,023 | 0,084 | 0,963 | 0,977 | 0,744 | 0 |
| 11 | 0,2 | 3 | 3 | 0,022 | 0,716 | 0,995 | 0,978 | 0,925 | 0,005 |
| 12 | 0,3 | 3 | 3 | 0,043 | 0,354 | 0,995 | 0,957 | 0,835 | 0,005 |
| 13 | 0,2 | 5 | 3 | 0,048 | 0,962 | 0,984 | 0,952 | 0,979 | 0,016 |
| 14 | 0,3 | 5 | 3 | 0,042 | 0,726 | 0,989 | 0,958 | 0,924 | 0,01 |
| 15 | 0,2 | 10 | 3 | 0,037 | 1 | 0,987 | 0,963 | 0,991 | 0,013 |
| 16 | 0,3 | 10 | 3 | 0,051 | 0,974 | 0,983 | 0,949 | 0,981 | 0,017 |
| 17 | 0,2 | 2 | 4 | 0,030 | 0,479 | 0,992 | 0,970 | 0,864 | 0,004 |
| 18 | 0,3 | 2 | 4 | 0,059 | 0,107 | 0,994 | 0,941 | 0,772 | 0,002 |
| 19 | 0,2 | 3 | 4 | 0,020 | 0,735 | 0,995 | 0,980 | 0,93 | 0,005 |
| 20 | 0,3 | 3 | 4 | 0,033 | 0,352 | 0,996 | 0,967 | 0,835 | 0,004 |
| 21 | 0,2 | 5 | 4 | 0,023 | 0,959 | 0,992 | 0,977 | 0,984 | 0,008 |
| 22 | 0,3 | 5 | 4 | 0,040 | 0,709 | 0,990 | 0,960 | 0,92 | 0,008 |
| 23 | 0,2 | 10 | 4 | 0,043 | 1 | 0,985 | 0,957 | 0,989 | 0,015 |
| 24 | 0,3 | 10 | 4 | 0,029 | 0,973 | 0,990 | 0,971 | 0,986 | 0,01 |

**Table S2:** Performance metrics for the PTM analysis of the 24 datasets in simulation 2. Essentially, the same conclusions can be drawn as from supplementary figure 2. MsqrobPTM generally performs very well, but performance can drop when the number of replicates gets too low, especially when combined with a higher variation or missingness present in the dataset.

#### Simulation 1 – peptidoform level

| SIMULATION 1 | | | | | | | | | |
|---|---|---|---|---|---|---|---|---|---|
| Dataset | sd | reps | conditions | fdp | sensitivity | specificity | precision | accuracy | fpr |
| 1 | 0,2 | 2 | 2 | 0,041 | 0,388 | 0,994 | 0,959 | 0,843 | 0,006 |
| 2 | 0,3 | 2 | 2 | 0,010 | 0,080 | 0,9997 | 0,990 | 0,770 | 0,0003 |
| 3 | 0,2 | 3 | 2 | 0,041 | 0,724 | 0,990 | 0,959 | 0,923 | 0,01 |
| 4 | 0,3 | 3 | 2 | 0,020 | 0,360 | 0,998 | 0,980 | 0,838 | 0,002 |
| 5 | 0,2 | 5 | 2 | 0,038 | 0,943 | 0,987 | 0,962 | 0,976 | 0,013 |
| 6 | 0,3 | 5 | 2 | 0,038 | 0,697 | 0,991 | 0,962 | 0,917 | 0,009 |
| 7 | 0,2 | 10 | 2 | 0,046 | 0,999 | 0,984 | 0,954 | 0,988 | 0,016 |
| 8 | 0,3 | 10 | 2 | 0,035 | 0,969 | 0,988 | 0,965 | 0,984 | 0,012 |
| 9 | 0,2 | 2 | 3 | 0,043 | 0,460 | 0,993 | 0,957 | 0,860 | 0,007 |
| 10 | 0,3 | 2 | 3 | 0,045 | 0,115 | 0,998 | 0,955 | 0,777 | 0,002 |
| 11 | 0,2 | 3 | 3 | 0,035 | 0,71 | 0,991 | 0,965 | 0,921 | 0,009 |
| 12 | 0,3 | 3 | 3 | 0,022 | 0,329 | 0,998 | 0,978 | 0,830 | 0,002 |
| 13 | 0,2 | 5 | 3 | 0,036 | 0,938 | 0,988 | 0,964 | 0,976 | 0,012 |
| 14 | 0,3 | 5 | 3 | 0,036 | 0,703 | 0,991 | 0,964 | 0,919 | 0,009 |
| 15 | 0,2 | 10 | 3 | 0,037 | 0,9998 | 0,987 | 0,963 | 0,990 | 0,013 |
| 16 | 0,3 | 10 | 3 | 0,048 | 0,976 | 0,983 | 0,952 | 0,982 | 0,017 |
| 17 | 0,2 | 2 | 4 | 0,030 | 0,413 | 0,996 | 0,970 | 0,850 | 0,004 |
| 18 | 0,3 | 2 | 4 | 0,038 | 0,098 | 0,999 | 0,962 | 0,774 | 0,001 |
| 19 | 0,2 | 3 | 4 | 0,035 | 0,711 | 0,991 | 0,965 | 0,921 | 0,009 |
| 20 | 0,3 | 3 | 4 | 0,041 | 0,364 | 0,995 | 0,959 | 0,837 | 0,005 |
| 21 | 0,2 | 5 | 4 | 0,042 | 0,947 | 0,986 | 0,958 | 0,976 | 0,013 |
| 22 | 0,3 | 5 | 4 | 0,029 | 0,723 | 0,993 | 0,971 | 0,925 | 0,007 |
| 23 | 0,2 | 10 | 4 | 0,035 | 0,9997 | 0,988 | 0,965 | 0,991 | 0,012 |
| 24 | 0,3 | 10 | 4 | 0,036 | 0,976 | 0,987 | 0,964 | 0,985 | 0,012 |

**Table S3:** Performance metrics for the peptidoform analysis of the 24 datasets in simulation 1. Essentially, the same conclusions can be drawn as from figure 3. FDR is well controlled on the peptidoform level as well, while still reporting a number of the significant peptidoforms. This number is high when there are many replicates, but can drop when the dataset is characterised by higher variation and a low number of repeats.

#### Simulation 2 – peptidoform level

| SIMULATION 2 | | | | | | | | | |
|---|---|---|---|---|---|---|---|---|---|
| Dataset | sd | reps | conditions | fdp | sensitivity | specificity | precision | accuracy | fpr |
| 1 | 0,2 | 2 | 2 | 0,021 | 0,242 | 0,5 | 0,979 | 0,437 | 0,002 |
| 2 | 0,3 | 2 | 2 | 0 | 0,072 | 0,509 | 1.000 | 0,402 | 0 |
| 3 | 0,2 | 3 | 2 | 0,023 | 0,431 | 0,910 | 0,977 | 0,791 | 0,003 |
| 4 | 0,3 | 3 | 2 | 0,056 | 0,171 | 0,910 | 0,944 | 0,726 | 0,003 |
| 5 | 0,2 | 5 | 2 | 0,041 | 0,832 | 0,986 | 0,959 | 0,948 | 0,012 |
| 6 | 0,3 | 5 | 2 | 0,022 | 0,534 | 0,994 | 0,978 | 0,879 | 0,004 |
| 7 | 0,2 | 10 | 2 | 0,027 | 1 | 0,991 | 0,973 | 0,993 | 0,009 |
| 8 | 0,3 | 10 | 2 | 0,049 | 0,926 | 0,984 | 0,951 | 0,970 | 0,016 |
| 9 | 0,2 | 2 | 3 | 0,055 | 0,194 | 0,646 | 0,945 | 0,534 | 0,004 |
| 10 | 0,3 | 2 | 3 | 0,029 | 0,069 | 0,673 | 0,971 | 0,523 | 0,0006 |
| 11 | 0,2 | 3 | 3 | 0,036 | 0,51 | 0,958 | 0,964 | 0,846 | 0,006 |
| 12 | 0,3 | 3 | 3 | 0,056 | 0,169 | 0,963 | 0,944 | 0,765 | 0,003 |
| 13 | 0,2 | 5 | 3 | 0,052 | 0,82 | 0,981 | 0,948 | 0,941 | 0,015 |
| 14 | 0,3 | 5 | 3 | 0,042 | 0,519 | 0,992 | 0,958 | 0,874 | 0,007 |
| 15 | 0,2 | 10 | 3 | 0,041 | 0,99 | 0,986 | 0,959 | 0,987 | 0,014 |
| 16 | 0,3 | 10 | 3 | 0,053 | 0,916 | 0,983 | 0,947 | 0,966 | 0,017 |
| 17 | 0,2 | 2 | 4 | 0,021 | 0,189 | 0,738 | 0,979 | 0,601 | 0,001 |
| 18 | 0,3 | 2 | 4 | 0,043 | 0,030 | 0,756 | 0,957 | 0,575 | 0,0004 |
| 19 | 0,2 | 3 | 4 | 0,019 | 0,446 | 0,963 | 0,981 | 0,834 | 0,002 |
| 20 | 0,3 | 3 | 4 | 0,027 | 0,218 | 0,969 | 0,973 | 0,781 | 0,002 |
| 21 | 0,2 | 5 | 4 | 0,026 | 0,837 | 0,992 | 0,974 | 0,953 | 0,008 |
| 22 | 0,3 | 5 | 4 | 0,044 | 0,507 | 0,991 | 0,956 | 0,870 | 0,008 |
| 23 | 0,2 | 10 | 4 | 0,036 | 0,993 | 0,988 | 0,964 | 0,989 | 0,012 |
| 24 | 0,3 | 10 | 4 | 0,033 | 0,924 | 0,990 | 0,967 | 0,973 | 0,01 |

**Table S4:** Performance metrics for the peptidoform analysis of the 24 datasets in simulation 2. Essentially, the same conclusions can be drawn as from figure 4. For the datasets with only 2 or 3 replicates the method starts to suffer from lack of information, making it harder to report any significant peptidoforms, especially for the datasets with sd 0.3. With higher amounts of replicates, the method still performs very well.

### Boxplots spike-in dataset

#### Boxplots showing effect of normalisation for PTM and protein dataset

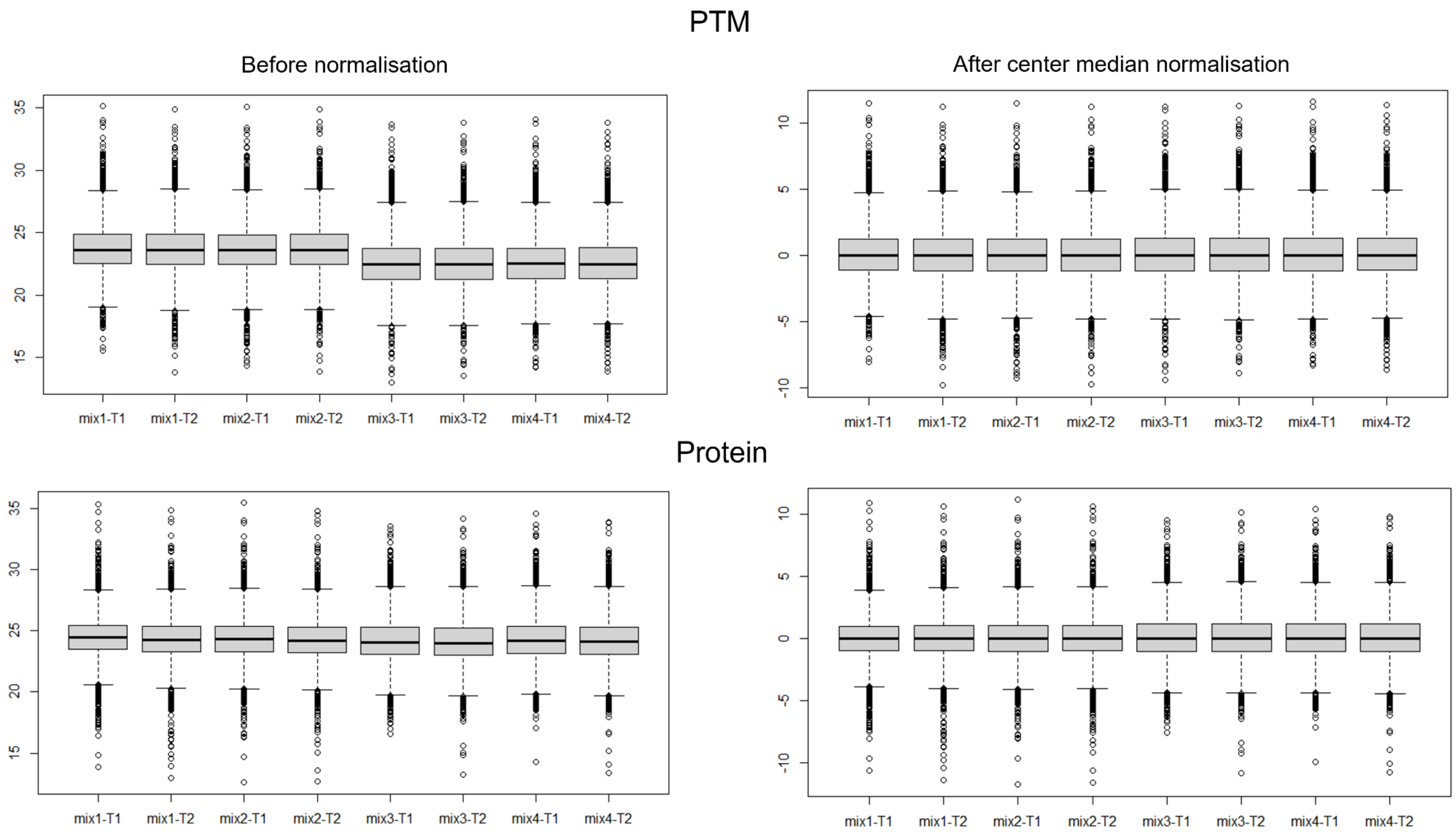

**Supplementary figure 3:** Boxplots of intensities for the spike-in experiment. At the top the intensities of the PTM data, at the bottom the intensities of the global proteome. The left plots show the intensities before median centring, the right plots after median centring. Before normalisation, at the PTM level mix 3 and 4 display a sudden drop in intensities in comparison to mix 1 and 2. This is because of the difference in background proteome used, with mix 3 and 4 containing a mix of E.coli and human proteins while mix 1 and 2 only contain the human proteins. After normalisation this drop disappears. Interestingly, this drop in intensities is not present at the protein level. Perhaps, this is the result of some pre-processing that was already done before the dataset was deposited on MASSIVE.

#### Tpr-fdp curves for each workflow and for each pairwise comparison

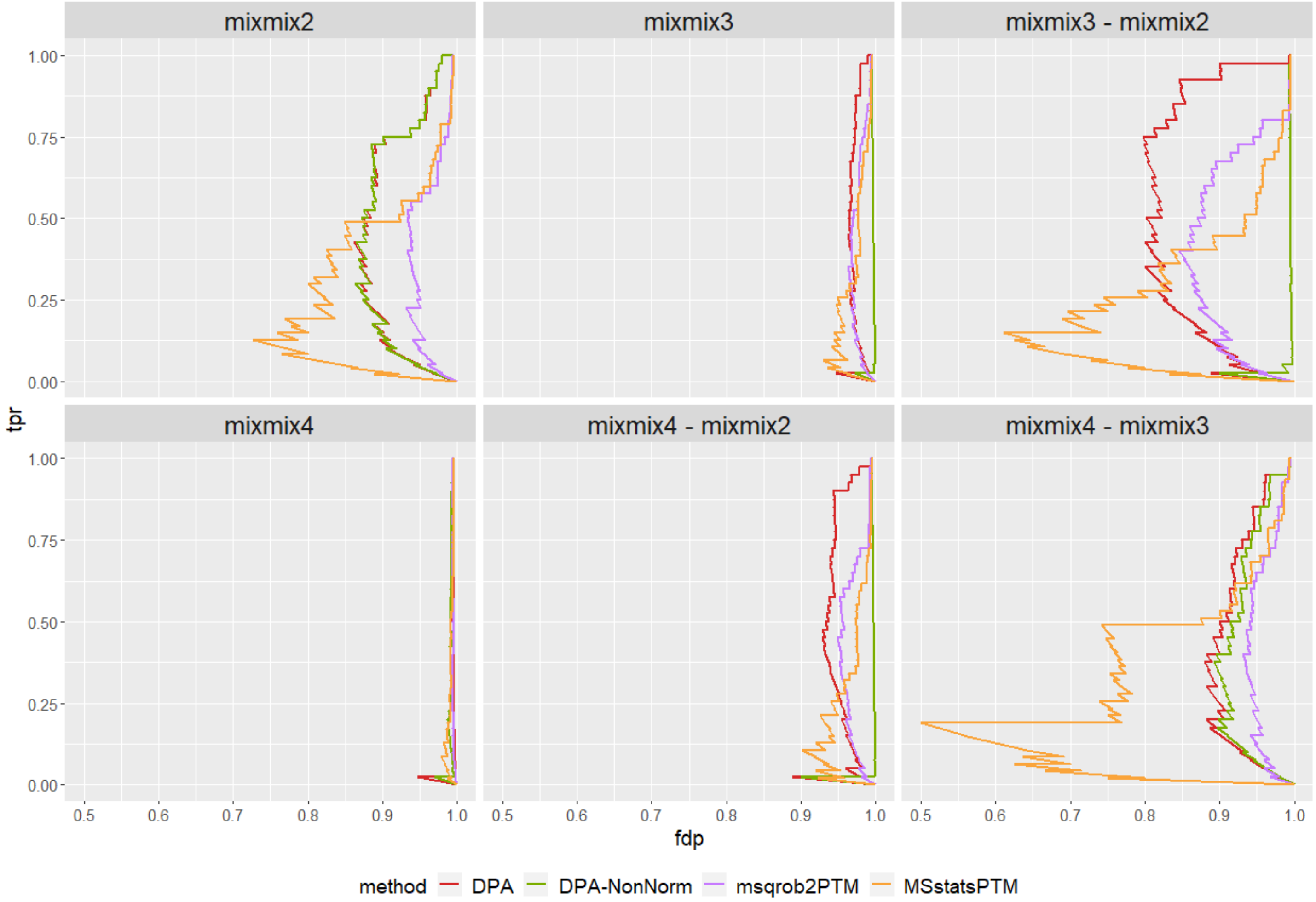

**Supplementary figure 4:** tpr-fdp curves for each workflow utilised on the spike-in dataset. DPA: differential PTM abundance by adopting a conventional msqrob2 workflow directly on the summarized PTM-level intensities, DPA-NonNorm: DPA but without normalisation with median peptidoform log-intensity, Msqrob2PTM: the standard msqrob2PTM workflow, MSstatsPTM: the standard MSstatsPTM workflow. Every square represents one pairwise comparison between the mixes.

#### Boxplots showing fold change of spike-in peptides

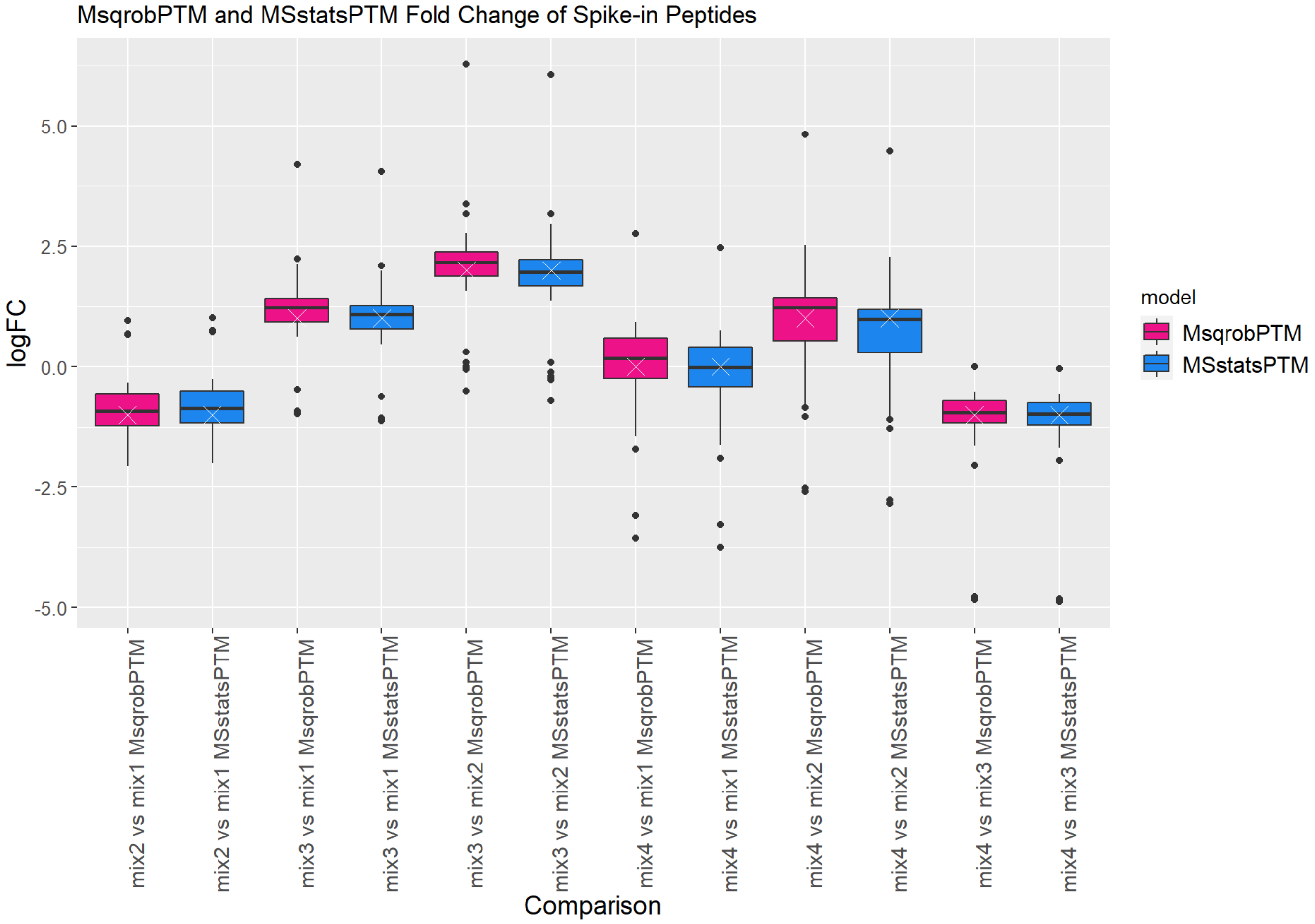

**Supplementary figure 5**: Boxplots show the distribution of the $log_2$ fold changes of the heavy peptides as estimated by the models, for both msqrobPTM (in pink) and MSstatsPTM (in blue). The white cross indicates the true fold change for a certain comparison of mixtures. The medians of the boxplots should coincide as well as possible with the crosses. Both models demonstrate good $log_2$ fold change estimation and very similar performance.

#### Boxplots showing fold change of background peptides

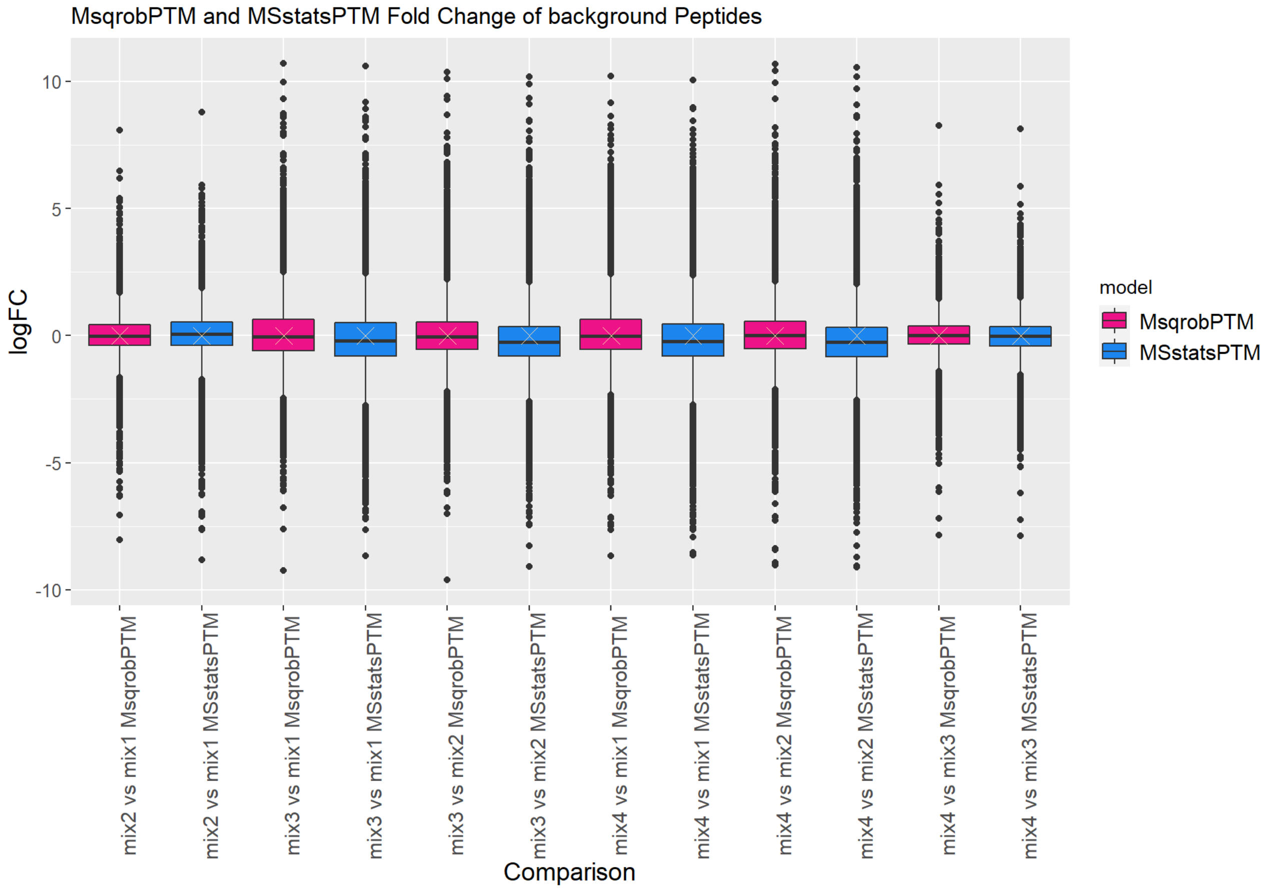

**Supplementary figure 6**: The boxplots show the distribution of the $\log_2$ fold changes of the background peptides as estimated by the models, for both msqrobPTM (in pink) and MSstatsPTM (in blue). The white cross indicates the true fold change for a certain comparison of mixtures, which is always zero for the light peptides as we do not expect them to change between mixtures. The medians of the boxplots should coincide as well as possible with the crosses. Both models show good $\log_2$ fold change estimation and very similar performance.

#### Detailed materials and methods for the biological phosphorylation dataset

##### Proteomics analysis

For the proteomic analysis, samples were precipitated with 100% MeOH in order to recover metabolites. After resuspension in 0.1% RapigestTM SF (Waters), in-solution digestion was performed on 20 µg proteins using Trypsin/Lys-C (Mass Spec Grade mix, Promega, Madison, USA) on an automated AssayMAP Bravo platform (Agilent Technologies). Samples were then incubated at 37°C for 45 minutes followed by centrifugation at 13 000 rpm for 10 minutes in order to remove Rapigest. Supernatants were collected and loaded on the AssayMAP Bravo platform (Agilent) to perform automated peptide clean-up on 5 µL phase C18 cartridges. Peptides were then resuspended in 150 µL of H2O and 0.1% FA.

NanoLC-MS/MS analyses were performed on a nanoAcquity UltraPerformance LC® (UPLC®) device (Waters Corporation, Milford, MA) coupled to a Q-Exactive Plus mass spectrometer (Thermo Fisher Scientific, Waltham, MA). The solvent system consisted of 0.1% FA in water (solvent A) and 0.1% FA in ACN (solvent B). Samples (equivalent of 400 ng of proteins) were loaded on a Symmetry C18 pre-column (20 mm × 180 µm with 5 µm diameter particles, Waters) over 3 min at 5 µL/min with 99% of solvent A and 1% of solvent B. Peptides were separated on an ACQUITY UPLC BEH130 C18 column (250 mm × 75 µm with 1.7 µm diameter particles) at 400 nL/min with the following gradient of solvent B: from 1 to 2 % over 2 min, from 2 to 25% over 77 min, from 25 to 35% over 10 minutes, from 35 to 90% over 1 minute then 90% for 5 minutes. Samples were injected in randomized order. The system was operated in DDA mode with automatic switching between MS (mass range 300–1800 m/z with R = 70,000, Automatic gain control (AGC) fixed at 3 x 106 ions and a maximum injection time set at 50 ms) and MS/MS (mass range 200–2000 m/z with R = 17,500, AGC fixed at 1 x 105 and the maximal injection time set to 100 ms) modes. The ten most abundant precursor ions were selected on each MS spectrum for further isolation and higher energy collision dissociation, excluding mono-charged and unassigned ions. The dynamic exclusion time was set to 60 s.

Raw data were processed using MaxQuant software (version 1.6.14). Peaks were assigned with the Andromeda search engine with trypsin/P specificity against an in-house generated protein sequence database containing all human entries extracted from UniProtKB-SwissProt (25th of August 2021, 20 339 entries). The minimal peptide length required was seven amino acids and a maximum of one missed cleavage was allowed. Methionine oxidation and acetylation of proteins' N-termini were set as variable modifications and Cysteine carbamidomethylation as a fixed modification. For protein quantification, the "match between runs" option was enabled. The maximum false discovery rate was set to 1% at peptide and protein levels with the use of a decoy strategy. Intensities were extracted from the Evidence.txt file to perform further statistical analysis.

##### Phosphoproteomics analysis

For the phosphoproteome analysis, protease inhibitors (Sigma, P8340) and phosphatase inhibitors (final concentration in Na3VO4 = 1 mM) were added to all samples. 100 µg proteins were used and total peptides extracts were prepared exactly as for total proteome analyses. The Phosphomix I light (Sigma Aldrich) was added to each sample (ratio peptide (µg)/mix(fmol) = 1.6) prior to phosphopeptide enrichment on 5 µL Fe(III)-NTA cartridges conducted on the AssayMAP Bravo platform following an IMAC protocol. After the enrichment, FA was added to each sample as well as phosphomix I heavy (Sigma Aldrich). Phosphopeptides were resuspended in 20 µL H2O, 2% ACN, 0.1% FA.

Nano-LC-MS/MS analyses were performed on a nanoAcquity UPLC devise (Waters) coupled to a Q-Exactive HF-X mass spectrometer (Thermo Scientific, Bremen, Germany) equipped with a Nanospray Flex™ ion source. The solvent system consisted of 0.1% FA in water (solvent A) and 0.1% FA in ACN (solvent B). Samples were loaded on an ACQUITY UPLC® Peptide BEH C18 Column (250 mm x 75 µm with 1.7 µm diameter particles) over 3 min at 5 µL/min with 99% of solvent A and 1% of solvent B. Phosphopeptides were separated on an ACQUITY UPLC® M-Class Symmetry® C18 Trap Column (20 mm x 180 µm with 5 µm diameter particles, Waters) at 350 nL/min with the following gradient of solvent B: from 1 to 2 % over 2 min, from 2 to 25% over 77 minutes, from 25 to 35% over 10 minutes and from 35 to 90% over 1 minute and finally 5 min at 90%. The system was operated in DDA mode with automatic switching between MS (mass range 375–1500 m/z with R = 120,000, AGC fixed at 3 x 106 ions and a maximum injection time set at 60 ms) and MS/MS (mass range 200–2000 m/z with R = 15,000, AGC fixed at 1 x 105 and the maximal injection time set to 60 ms) modes. The ten most abundant ions were selected on each MS spectrum for further isolation and higher energy collision dissociation, excluding mono-charged and unassigned ions. The dynamic exclusion time was set to 40 s.

Raw data files were processed using MaxQuant software (version 1.6.14). Peaks were assigned with the Andromeda search engine with trypsin/P specificity against an in-house generated protein sequence database containing all human entries extracted from UniProtKB-SwissProt (25th of August 2021, 20 339 entries). The minimal peptide length required was seven amino acids and a maximum of one missed cleavage was allowed. Methionine oxidation, acetylation of proteins' N-termini and serine, threonine and tyrosine phosphorylation were set as variable modifications and Cysteine carbamidomethylation as a fixed modification. For protein quantification, the "match between runs" option was enabled. The maximum false discovery rate was set to 1% at peptide and protein levels with the use of a decoy strategy. Intensities were extracted from the Evidence.txt file to perform the further statistical analysis.

### Detailed results of the phosphorylation dataset

#### Workflow with only the enriched data

| PEPTIDOFORM | | | | | | | | |
|---|---|---|---|---|---|---|---|---|
| **contrast** | **peptidoform** | **logFC** | **se** | **df** | **t** | **pval** | **adjPval** | **rank** |
| conditionA | VGYVSGWGR | -1.447473 | 0.3 | 37 | -4.8 | 2.98e-05 | 0.0463 | 1 |
| conditionA + conditionA:subsety | VS(Phospho (STY))REFHSHEFHSHEDM(Oxidation (M))LVVDPK | -1.314907 | 0.30 | 71.6 | -4.3 | 0.0000450 | 0.0361 | 1 |
| conditionA + conditionA:subsety | EPQDTYHYLPFS(Phospho (STY))LPHR | -1.656762 | 0.36 | 34.6 | -4.6 | 0.0000563 | 0.0361 | 2 |
| conditionA + conditionA:subsety | LPIVNFDYS(Phospho (STY))M(Oxidation (M))EEK | -0.781947 | 0.19 | 75.6 | -4.2 | 0.0000697 | 0.0361 | 3 |
| conditionA + conditionA:subsety | LNVEDVDSTK | 0.6265500 | 0.15 | 49.6 | 4.2 | 0.0001015 | 0.0395 | 4 |
| conditionA + conditionA:subsety | AAM(Oxidation (M))VGMLANFLGFR | -3.128131 | 0.19 | 3.6 | -16.4 | 0.0001530 | 0.0476 | 5 |
| conditionA + conditionA:subsety | ADQTVLTEDEK | 1.0553334 | 0.27 | 70.6 | 3.9 | 0.0001927 | 0.0500 | 6 |
| conditionA + 0.5 * conditionA:subsety | VGYVSGWGR | -0.815019 | 0.17 | 37 | -4.9 | 2.01e-05 | 0.0313 | 1 |
| conditionA + 0.6666667 * conditionA:subsety | LPIVNFDYS(Phospho (STY))M(Oxidation (M))EEK | -0.708945 | 0.15 | 75.6 | -4.6 | 0.0000153 | 0.0238 | 1 |
| conditionA + 0.6666667 * conditionA:subsety | VGYVSGWGR | -0.604200 | 0.14 | 36.6 | -4.4 | 0.0000812 | 0.0494 | 2 |
| conditionA + 0.6666667 * conditionA:subsety | VS(Phospho (STY))REFHSHEFHSHEDM(Oxidation (M))LVVDPK | -0.990202 | 0.24 | 71.6 | -4.1 | 0.0001164 | 0.0494 | 3 |
| conditionA + 0.6666667 * conditionA:subsety | AAM(Oxidation (M))VGMLANFLGFR | -2.485975 | 0.14 | 3.6 | -17.3 | 0.0001270 | 0.0494 | 4 |
| **PTM** | | | | | | | | |
| **contrast** | **ptm** | **logFC** | **se** | **df** | **t** | **pval** | **adjPval** | **rank** |
| conditionA + conditionA:subsety | sp\|P01019\|ANGT_HUMAN (Oxidation (M)) 105 | -3.288795 | 0.16 | 4.8 | -20.8 | 0.0000068 | 0.00193 | 1 |
| conditionA + conditionA:subsety | sp\|P10909\|CLUS_HUMAN (Phospho (STY)) 210 | -1.673530 | 0.33 | 32.9 | -5.1 | 0.0000128 | 0.00193 | 2 |

| | | | | | | | | |
|---|---|---|---|---|---|---|---|---|
| conditionA + conditionA:subsety | sp\|P10451\|OSTP_HUMAN (Oxidation (M)) 284 | -0.765036 | 0.17 | 82.4 | -4.6 | 0.0000144 | 0.00193 | 3 |
| conditionA + conditionA:subsety | sp\|P02765\|FETUA_HUMAN (Oxidation (M)) 321 | -1.217209 | 0.29 | 68.9 | -4.1 | 0.0000962 | 0.00969 | 4 |
| conditionA + conditionA:subsety | sp\|O94769\|ECM2_HUMAN (Oxidation (M)) 76 | -0.694987 | 0.18 | 73.2 | -3.9 | 0.0001927 | 0.01366 | 5 |
| conditionA + conditionA:subsety | sp\|P05060\|SCG1_HUMAN (Phospho (STY)) 317 | -0.835026 | 0.21 | 62.4 | -3.9 | 0.0002034 | 0.01366 | 6 |
| conditionA + conditionA:subsety | sp\|O94769\|ECM2_HUMAN (Phospho (STY)) 75 | -0.730700 | 0.19 | 80.5 | -3.8 | 0.0002810 | 0.01607 | 7 |
| conditionA + conditionA:subsety | sp\|P05060\|SCG1_HUMAN (Phospho (STY)) 311 | -0.770509 | 0.20 | 81.8 | -3.8 | 0.0003190 | 0.01607 | 8 |
| conditionA + conditionA:subsety | sp\|P62328\|TYB4_HUMAN (Acetyl (Protein N-term)) 1 | 0.8411341 | 0.18 | 13.0 | 4.8 | 0.0003640 | 0.01630 | 9 |
| conditionA + conditionA:subsety | sp\|O94769\|ECM2_HUMAN (Phospho (STY)) 245 | 0.4363289 | 0.12 | 68.0 | 3.6 | 0.0005450 | 0.02155 | 10 |
| conditionA + conditionA:subsety | sp\|P01042\|KNG1_HUMAN (Phospho (STY)) 275 | -1.32233 | 0.35 | 33.2 | -3.8 | 0.0005882 | 0.02155 | 11 |
| conditionA + conditionA:subsety | sp\|P10451\|OSTP_HUMAN (Phospho (STY)) 280 | -1.075666 | 0.31 | 84.1 | -3.5 | 0.0006994 | 0.02349 | 12 |
| conditionA:subsety | sp\|P10909\|CLUS_HUMAN (Phospho (STY)) 210 | -2.363970 | 0.52 | 32.9 | -4.6 | 0.0000680 | 0.021 | 1 |
| conditionA:subsety | sp\|P01019\|ANGT_HUMAN (Oxidation (M)) 105 | -2.440847 | 0.21 | 4.8 | -11.7 | 0.0001041 | 0.021 | 2 |
| conditionA + 0.5 * conditionA:subsety | sp\|P01019\|ANGT_HUMAN (Oxidation (M)) 105 | -2.068372 | 0.10 | 4.8 | -19.8 | 0.0000086 | 0.00349 | 1 |
| conditionA + 0.5 * conditionA:subsety | sp\|O94769\|ECM2_HUMAN (Oxidation (M)) 76 | -0.601516 | 0.16 | 73.2 | -3.9 | 0.0002451 | 0.03315 | 2 |
| conditionA + 0.5 * conditionA:subsety | sp\|P07197\|NFM_HUMAN (Phospho (STY)) 736 | -1.425155 | 0.33 | 23.6 | -4.3 | 0.0002739 | 0.03315 | 3 |
| conditionA + 0.5 * conditionA:subsety | sp\|Q14515\|SPRL1_HUMAN (Oxidation (M)) 276 | -1.563138 | 0.38 | 29.1 | -4.1 | 0.0003290 | 0.03315 | 4 |
| conditionA + 0.6666667 * conditionA:subsety | sp\|P01019\|ANGT_HUMAN (Oxidation (M)) 105 | -2.47518 | 0.11 | 4.8 | -21.6 | 0.0000058 | 0.00232 | 1 |
| conditionA + 0.6666667 * conditionA:subsety | sp\|O94769\|ECM2_HUMAN (Oxidation (M)) 76 | -0.632673 | 0.15 | 73.2 | -4.3 | 0.0000450 | 0.00906 | 2 |

| | | | | | | | | |
|---|---|---|---|---|---|---|---|---|
| conditionA + 0.6666667 * conditionA:subsety | sp\|P07197\|NFM_HUMAN (Phospho (STY)) 736 | -1.416621 | 0.32 | 23.6 | -4.4 | 0.0001969 | 0.02640 | 3 |
| conditionA + 0.6666667 * conditionA:subsety | sp\|P10451\|OSTP_HUMAN (Oxidation (M)) 284 | -0.516539 | 0.14 | 82.4 | -3.8 | 0.0002620 | 0.02640 | 4 |
| conditionA + 0.6666667 * conditionA:subsety | sp\|Q14515\|SPRL1_HUMAN (Oxidation (M)) 276 | -1.481238 | 0.38 | 29.1 | -3.9 | 0.0005106 | 0.03572 | 5 |
| conditionA + 0.6666667 * conditionA:subsety | sp\|O94769\|ECM2_HUMAN (Phospho (STY)) 75 | -0.556072 | 0.16 | 80.5 | -3.6 | 0.0006166 | 0.03572 | 6 |
| conditionA + 0.6666667 * conditionA:subsety | sp\|P02765\|FETUA_HUMAN (Oxidation (M)) 321 | -0.873004 | 0.24 | 68.9 | -3.6 | 0.0006205 | 0.03572 | 7 |
| conditionA + 0.6666667 * conditionA:subsety | sp\|P01042\|KNG1_HUMAN (Phospho (STY)) 275 | -1.020457 | 0.28 | 33.2 | -3.7 | 0.0008739 | 0.04240 | 8 |
| conditionA + 0.6666667 * conditionA:subsety | sp\|P05060\|SCG1_HUMAN (Phospho (STY)) 311 | -0.569902 | 0.17 | 81.8 | -3.4 | 0.0009469 | 0.04240 | 9 |

**Table S5:** all significant peptidoforms and PTMs for the biological phospho dataset when using only the enriched data. The data was modeled with the variables indicated in the metadata. The researchers wanted to know whether there is a difference in PTM usage between condition A and condition B, and whether that difference is different for samples from different subsets (x or y). We can specify this model by using a formula with the factor condition and subset as its predictors: formula = ~condition*subset. Note that we include the interaction effect by using an asterisk in the formula. Also note that a formula always starts with a tilde '~'. When using this model, the following coefficients are calculated: (Intercept), conditionA, subsety and conditionA:subsety (interaction). Condition B is the reference class for condition and subset x is the reference class for subset. So the mean log2 expression for B and x samples is '(Intercept)'. The mean log2 expression for A and x samples is '(Intercept)+ conditionA. Hence, the average log2 fold change between condition A and B for subset x is modeled using the parameter 'conditionA'. Thus, we assess the contrast 'conditionA=0' with our statistical test. In the same way, the average log2 fold change between condition A and B for subset y is modeled using the parameter 'conditionA + conditionA:subsety'. Thus, we assess the contrast 'conditionA + conditionA:subsety = 0' with our statistical test to find PTMs differential between condition A and B for subset y.
The difference between abovementioned contrasts is modelled using the interaction parameter: conditionA:subsety. When we assess this contrast, we will obtain PTMS that exhibit different changes in subset x between condition A and B, as compared to subset y between condition A and B.
We can also assess an average contrast to see the average difference between A and B samples across subsets: conditionA + 0.5 * conditionA:subsety = 0 and the marginal effect of condition: conditionA + 0.6666667 * conditionA:subsety (with 0.6666667 = number of y samples / total number of samples).

#### Workflow using both datasets

| PEPTIDOFORM | | | | | | | | |
|---|---|---|---|---|---|---|---|---|
| **contrast** | **peptidoform** | **logFC** | **se** | **df** | **t** | **pval** | **adjPval** | **rank** |
| conditionA + conditionA:subsety | LVGGPM(Oxidation (M))DAS(Phospho (STY))VEEEGVRR | -1.254296 | 0.25 | 87 | -5.1 | 0.0000023 | 0.00249 | 1 |
| conditionA + conditionA:subsety | VS(Phospho (STY))REFHSHEFHSHEDM(Oxidation (M))LVVDPK | -2.016171 | 0.40 | 76 | -5.0 | 0.0000035 | 0.00249 | 2 |
| conditionA + conditionA:subsety | EFHSHEFHS(Phospho (STY))HEDM(Oxidation (M))LVVDPK | -1.86826 | 0.41 | 87 | -4.5 | 0.0000194 | 0.00913 | 3 |
| conditionA + conditionA:subsety | LPIVNFDYS(Phospho (STY))M(Oxidation (M))EEK | -1.061565 | 0.24 | 80 | -4.4 | 0.0000393 | 0.01390 | 4 |
| conditionA + conditionA:subsety | GHPQEESEESNVS(Phospho (STY))MASLGEK | -1.173063 | 0.27 | 49 | -4.4 | 0.0000618 | 0.01748 | 5 |
| conditionA + conditionA:subsety | LVGGPMDAS(Phospho (STY))VEEEGVRR | -1.441447 | 0.35 | 90 | -4.1 | 0.0000965 | 0.02274 | 6 |
| conditionA + conditionA:subsety | S(Phospho (STY))GEATDGARPQALPEPMQESK | -1.35189 | 0.34 | 85 | -4.0 | 0.0001483 | 0.02798 | 7 |
| conditionA + conditionA:subsety | EIPAWVPFDPAAQITK | -2.016909 | 0.45 | 25 | -4.4 | 0.0001583 | 0.02798 | 8 |
| conditionA + conditionA:subsety | EFHS(Phospho (STY))HEFHS(Phospho (STY))HEDM(Oxidation (M))LVVDPK | -1.694843 | 0.44 | 81 | -3.9 | 0.0002024 | 0.03180 | 9 |
| conditionA + conditionA:subsety | GILAADESTGS(Phospho (STY))IAKR | -0.712955 | 0.19 | 81 | -3.8 | 0.0002476 | 0.03344 | 10 |
| conditionA + conditionA:subsety | EFHSHEFHS(Phospho (STY))HEDMLVVDPK | -2.153129 | 0.56 | 81 | -3.8 | 0.0002602 | 0.03344 | 11 |
| conditionA + 0.6666667 * conditionA:subsety | LVGGPM(Oxidation (M))DAS(Phospho (STY))VEEEGVRR | -0.920206 | 0.20 | 87 | -4.6 | 0.0000136 | 0.0193 | 1 |
| conditionA + 0.6666667 * conditionA:subsety | VS(Phospho (STY))REFHSHEFHSHEDM(Oxidation (M))LVVDPK | -1.42027 | 0.32 | 76 | -4.4 | 0.0000358 | 0.0220 | 2 |
| conditionA + 0.6666667 * conditionA:subsety | LPIVNFDYS(Phospho (STY))M(Oxidation (M))EEK | -0.867057 | 0.20 | 80 | -4.3 | 0.0000466 | 0.0220 | 3 |
| conditionA + 0.6666667 * conditionA:subsety | GHPQEESEESNVS(Phospho (STY))MASLGEK | -0.950535 | 0.22 | 49 | -4.3 | 0.0000690 | 0.0244 | 4 |
| conditionA + 0.6666667 * conditionA:subsety | GILAADESTGS(Phospho (STY))IAK | -0.722465 | 0.18 | 73 | -4.0 | 0.0001662 | 0.0470 | 5 |
| **PTM** | | | | | | | | |
| **contrast** | **ptm** | **logFC** | **se** | **df** | **t** | **pval** | **adjPval** | **rank** |
| conditionA + conditionA:subsety | sp\|P10451\|OSTP_HUMAN (Oxidation (M)) 284 | -1.465966 | 0.31 | 89 | -4.8 | 0.0000075 | 0.00279 | 1 |
| conditionA + conditionA:subsety | sp\|P01034\|CYTC_HUMAN (Oxidation (M)) 40 | -0.971481 | 0.22 | 88 | -4.3 | 0.0000393 | 0.00509 | 2 |
| conditionA + conditionA:subsety | sp\|O94769\|ECM2_HUMAN (Oxidation (M)) 76 | -1.061565 | 0.24 | 79 | -4.3 | 0.0000413 | 0.00509 | 3 |
| conditionA + conditionA:subsety | sp\|P10645\|CMGA_HUMAN (Phospho (STY)) 161 | -1.278335 | 0.30 | 61 | -4.2 | 0.0000795 | 0.00735 | 4 |

| | | | | | | | | |
|---|---|---|---|---|---|---|---|---|
| conditionA + conditionA:subsety | sp\|P10645\|CMGA_HUMAN (Phospho (STY)) 142 | -1.257455 | 0.31 | 84 | -4.1 | 0.0001125 | 0.00753 | 5 |
| conditionA + conditionA:subsety | sp\|P04075\|ALDOA_HUMAN (Phospho (STY)) 39 | -0.710081 | 0.18 | 88 | -4.0 | 0.0001222 | 0.00753 | 6 |
| conditionA + conditionA:subsety | sp\|P05060\|SCG1_HUMAN (Phospho (STY)) 317 | -1.138593 | 0.30 | 70 | -3.8 | 0.0002791 | 0.01475 | 7 |
| conditionA + conditionA:subsety | sp\|P10451\|OSTP_HUMAN (Phospho (STY)) 280 | -1.789866 | 0.50 | 89 | -3.6 | 0.0005233 | 0.02275 | 8 |
| conditionA + conditionA:subsety | sp\|O94769\|ECM2_HUMAN (Phospho (STY)) 75 | -1.100803 | 0.31 | 89 | -3.6 | 0.0005534 | 0.02275 | 9 |
| conditionA + conditionA:subsety | sp\|P02774\|VTDB_HUMAN (Phospho (STY)) 95 | -0.771993 | 0.22 | 63 | -3.6 | 0.0006759 | 0.02501 | 10 |
| conditionA + conditionA:subsety | sp\|P01042\|KNG1_HUMAN (Phospho (STY)) 275 | -1.353078 | 0.38 | 39 | -3.6 | 0.0009266 | 0.02893 | 11 |
| conditionA + conditionA:subsety | sp\|P51693\|APLP1_HUMAN (Phospho (STY)) 515 | -1.295304 | 0.38 | 83 | -3.4 | 0.0009384 | 0.02893 | 12 |
| conditionA + conditionA:subsety | sp\|P10451\|OSTP_HUMAN (Phospho (STY)) 27 | -1.254945 | 0.37 | 89 | -3.4 | 0.0010416 | 0.02964 | 13 |
| conditionA + conditionA:subsety | sp\|P10645\|CMGA_HUMAN (Phospho (STY)) 218 | -0.914380 | 0.27 | 89 | -3.4 | 0.0011272 | 0.02978 | 14 |
| conditionA + conditionA:subsety | sp\|P13521\|SCG2_HUMAN (Phospho (STY)) 106 | -1.237754 | 0.36 | 48 | -3.4 | 0.0012072 | 0.02978 | 15 |
| conditionA + conditionA:subsety | sp\|P02765\|FETUA_HUMAN (Oxidation (M)) 321 | -1.443326 | 0.45 | 79 | -3.2 | 0.0017541 | 0.03823 | 16 |
| conditionA + conditionA:subsety | sp\|P09972\|ALDOC_HUMAN (Phospho (STY)) 39 | -1.168275 | 0.36 | 73 | -3.2 | 0.0019930 | 0.03823 | 17 |
| conditionA + conditionA:subsety | sp\|P19823\|ITIH2_HUMAN (Oxidation (M)) 64 | -2.462465 | 0.69 | 19 | -3.6 | 0.0020215 | 0.03823 | 18 |
| conditionA + conditionA:subsety | sp\|P10451\|OSTP_HUMAN (Phospho (STY)) 24 | -0.966383 | 0.30 | 62 | -3.2 | 0.0021278 | 0.03823 | 19 |
| conditionA + conditionA:subsety | sp\|P10909\|CLUS_HUMAN (Phospho (STY)) 210 | -1.369371 | 0.42 | 38 | -3.3 | 0.0021492 | 0.03823 | 20 |
| conditionA + conditionA:subsety | sp\|P00747\|PLMN_HUMAN (Phospho (STY)) 358 | -0.978169 | 0.30 | 58 | -3.2 | 0.0021699 | 0.03823 | 21 |
| conditionA + conditionA:subsety | sp\|P24592\|IBP6_HUMAN (Phospho (STY)) 152 | -0.860173 | 0.28 | 86 | -3.1 | 0.0025077 | 0.04098 | 22 |
| conditionA + conditionA:subsety | sp\|P10451\|OSTP_HUMAN (Phospho (STY)) 195 | -1.246640 | 0.40 | 76 | -3.1 | 0.0025475 | 0.04098 | 23 |
| conditionA + conditionA:subsety | sp\|P01034\|CYTC_HUMAN (Phospho (STY)) 43 | -0.960092 | 0.31 | 90 | -3.1 | 0.0027547 | 0.04247 | 24 |
| conditionA + conditionA:subsety | sp\|P24593\|IBP5_HUMAN (Phospho (STY)) 124 | -0.883168 | 0.29 | 66 | -3.1 | 0.0029382 | 0.04348 | 25 |
| conditionA + conditionA:subsety | sp\|P10645\|CMGA_HUMAN (Phospho (STY)) 370 | -0.917409 | 0.29 | 35 | -3.2 | 0.0032049 | 0.04561 | 26 |
| conditionA + conditionA:subsety | sp\|P61769\|B2MG_HUMAN (Phospho (STY)) 108 | -0.60206 | 0.20 | 87 | -3.0 | 0.0034231 | 0.04592 | 27 |
| conditionA + conditionA:subsety | sp\|P30086\|PEBP1_HUMAN (Phospho (STY)) 52 | -0.990936 | 0.33 | 80 | -3.0 | 0.0034749 | 0.04592 | 28 |
| conditionA + conditionA:subsety | sp\|P04075\|ALDOA_HUMAN (Phospho (STY)) 36 | -1.209235 | 0.40 | 79 | -3.0 | 0.0037198 | 0.04746 | 29 |
| conditionA + conditionA:subsety | sp\|P05060\|SCG1_HUMAN (Phospho (STY)) 311 | -0.860726 | 0.29 | 90 | -3.0 | 0.0039602 | 0.04884 | 30 |
| conditionA + 0.6666667 * conditionA:subsety | sp\|P04075\|ALDOA_HUMAN (Phospho (STY)) 39 | -0.642709 | 0.14 | 88 | -4.5 | 0.0000246 | 0.00905 | 1 |
| conditionA + 0.6666667 * conditionA:subsety | sp\|O94769\|ECM2_HUMAN (Oxidation (M)) 76 | -0.867057 | 0.20 | 79 | -4.3 | 0.0000489 | 0.00905 | 2 |

| | | | | | | | | |
|---|---|---|---|---|---|---|---|---|
| conditionA + 0.6666667 * conditionA:subsety | sp\|P01034\|CYTC_HUMAN (Oxidation (M)) 40 | -0.716407 | 0.18 | 88 | -3.9 | 0.0001623 | 0.02002 | 3 |
| conditionA + 0.6666667 * conditionA:subsety | sp\|P10451\|OSTP_HUMAN (Oxidation (M)) 284 | -0.963708 | 0.25 | 89 | -3.8 | 0.0002268 | 0.02098 | 4 |
| conditionA + 0.6666667 * conditionA:subsety | sp\|P10645\|CMGA_HUMAN (Phospho (STY)) 142 | -0.926145 | 0.25 | 84 | -3.7 | 0.0003919 | 0.02900 | 5 |
| conditionA + 0.6666667 * conditionA:subsety | sp\|P02774\|VTDB_HUMAN (Phospho (STY)) 95 | -0.64878 | 0.18 | 63 | -3.7 | 0.0005014 | 0.03092 | 6 |
| conditionA + 0.6666667 * conditionA:subsety | sp\|P30086\|PEBP1_HUMAN (Phospho (STY)) 52 | -0.940872 | 0.26 | 80 | -3.6 | 0.0006198 | 0.03276 | 7 |
| conditionA + 0.6666667 * conditionA:subsety | sp\|P09972\|ALDOC_HUMAN (Phospho (STY)) 39 | -1.000408 | 0.29 | 73 | -3.5 | 0.0008802 | 0.04071 | 8 |
| conditionA + 0.6666667 * conditionA:subsety | sp\|Q14515\|SPRL1_HUMAN (Oxidation (M)) 276 | -1.530703 | 0.43 | 35 | -3.5 | 0.0012155 | 0.04822 | 9 |
| conditionA + 0.6666667 * conditionA:subsety | sp\|P10645\|CMGA_HUMAN (Phospho (STY)) 218 | -0.733797 | 0.22 | 89 | -3.3 | 0.0013032 | 0.04822 | 10 |
| conditionA + 0.6666667 * conditionA:subsety | sp\|P10645\|CMGA_HUMAN (Phospho (STY)) 161 | -0.836548 | 0.25 | 61 | -3.3 | 0.0014779 | 0.04971 | 11 |
| conditionA + 0.6666667 * conditionA:subsety | sp\|P04075\|ALDOA_HUMAN (Phospho (STY)) 36 | -1.079675 | 0.33 | 79 | -3.3 | 0.0016171 | 0.04986 | 12 |

**Table S6: all significant peptidoforms and PTMs for the biological phospho dataset when using both the enriched and non-enriched data**

### Additional examples of the mock analysis

#### Workflow with only enriched dataset

##### *Example 1*

PTM level

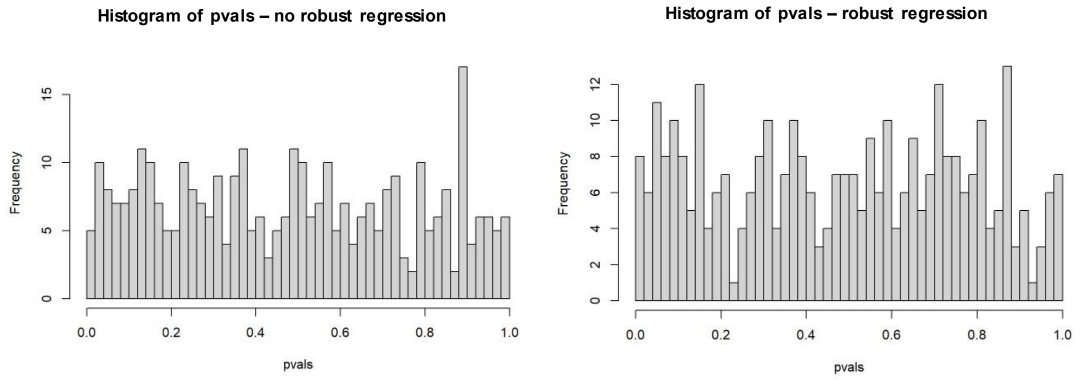

Peptidoform level

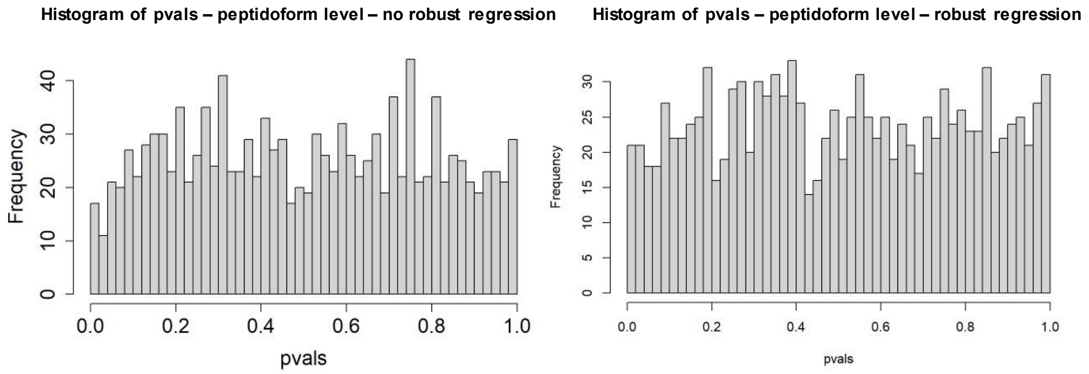

**Supplementary figure 7:** Distribution of p-values for the mock analysis of the phospho dataset without the use of a global profiling run, for analysis on PTM level (top) as well as peptidoform level (bottom). For the plots on the left, the analysis was carried out without robust regression during the modelling step. In contrast, the plots on the right correspond to an analysis carried out with robust regression during the modelling step.

#### *Example 2*

PTM level

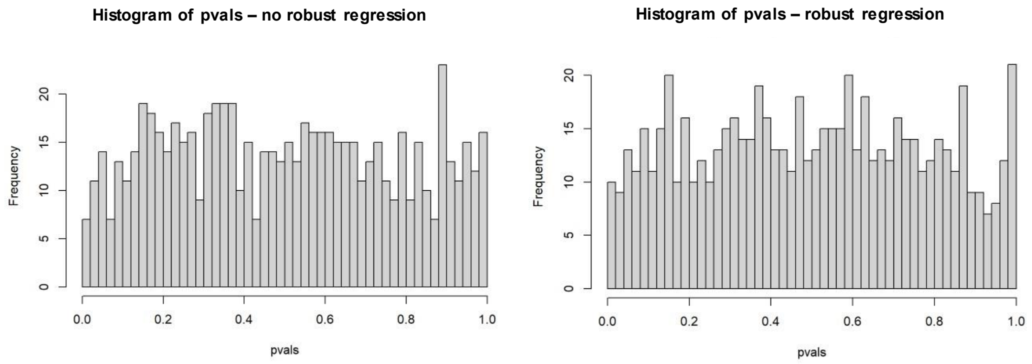

Peptidoform level

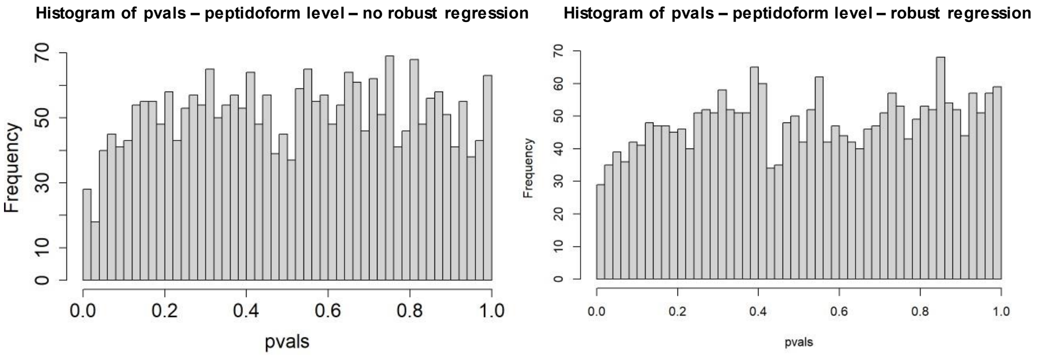

**Supplementary figure 8:** Distribution of p-values for the mock analysis of the phospho dataset without the use of a global profiling run, for analysis on PTM level (top) as well as peptidoform level (bottom). For the plots on the left, the analysis was carried out without robust regression during the modelling step. In contrast, the plots on the right correspond to an analysis carried out with robust regression during the modelling step.

#### *Example 3*

PTM level

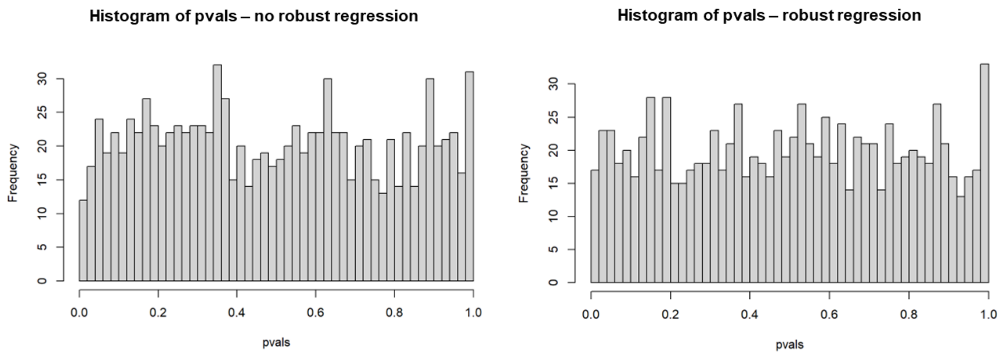

Peptidoform level

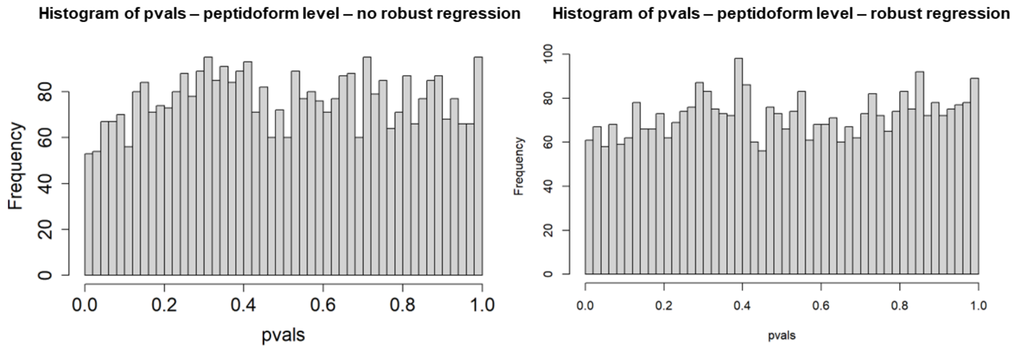

**Supplementary figure 9:** Distribution of p-values for the mock analysis of the phospho dataset without the use of a global profiling run, for analysis on PTM level (top) as well as peptidoform level (bottom). For the plots on the left, the analysis was carried out without robust regression during the modelling step. In contrast, the plots on the right correspond to an analysis carried out with robust regression during the modelling step.

#### *Example 4*

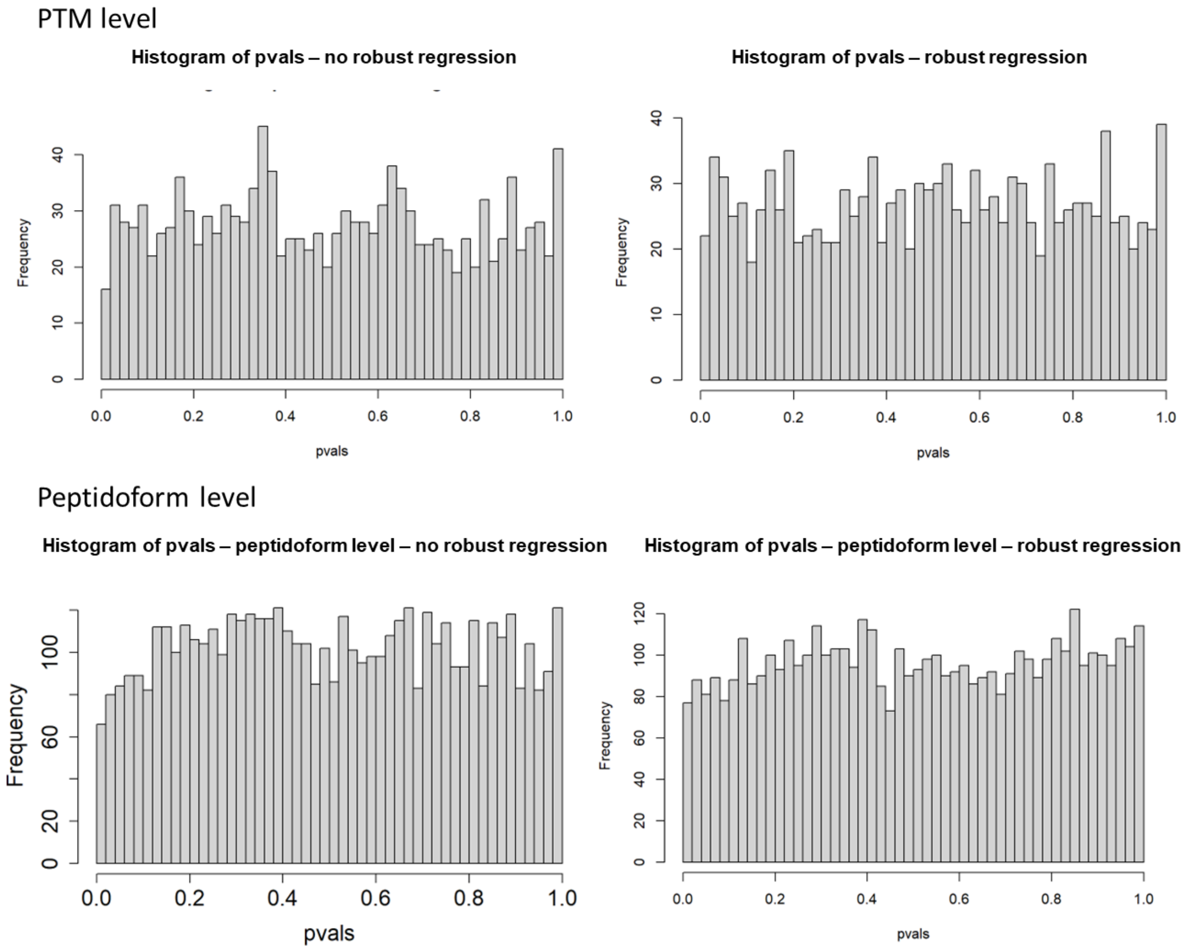

**Supplementary figure 10:** Distribution of p-values for the mock analysis of the phospho dataset without the use of a global profiling run, for analysis on PTM level (top) as well as peptidoform level (bottom). For the plots on the left, the analysis was carried out without robust regression during the modelling step. In contrast, the plots on the right correspond to an analysis carried out with robust regression during the modelling step.

#### Workflow including non-enriched counterpart

##### *Example 1*

PTM level

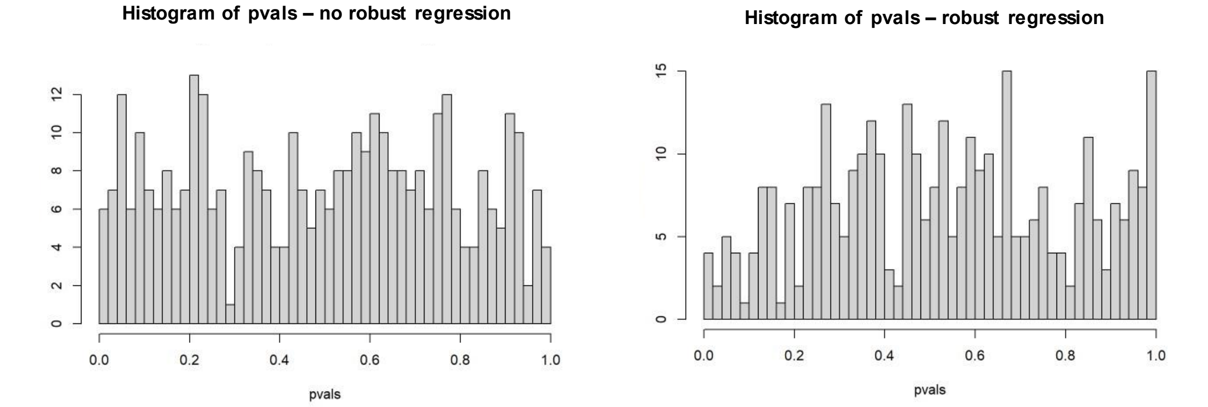

Peptidoform level

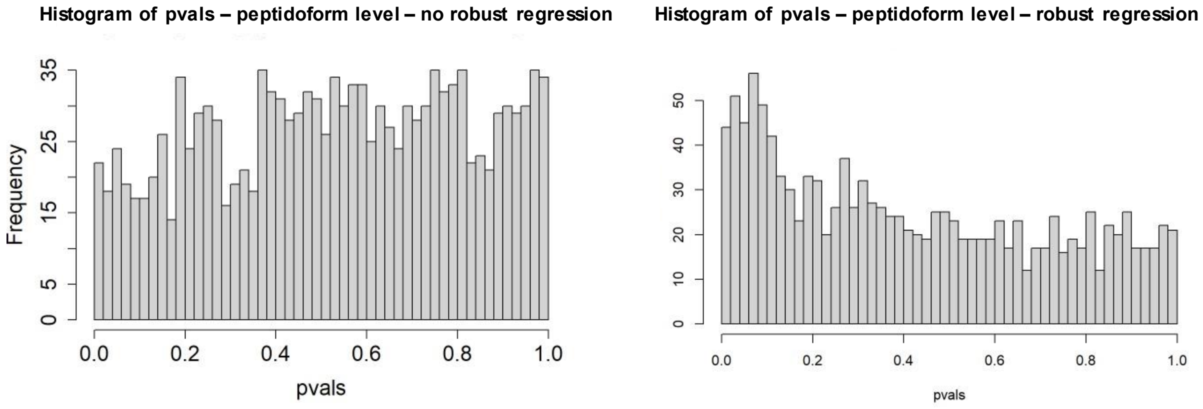

**Supplementary figure 11:** Distribution of p-values for the mock analysis of the phospho dataset including the use of a global profiling run, for analysis on PTM level (top) as well as peptidoform level (bottom). For the plots on the left, the analysis was carried out without robust regression during the modelling step. In contrast, the plots on the right correspond to an analysis carried out with robust regression during the modelling step.

#### *Example 2*

PTM level

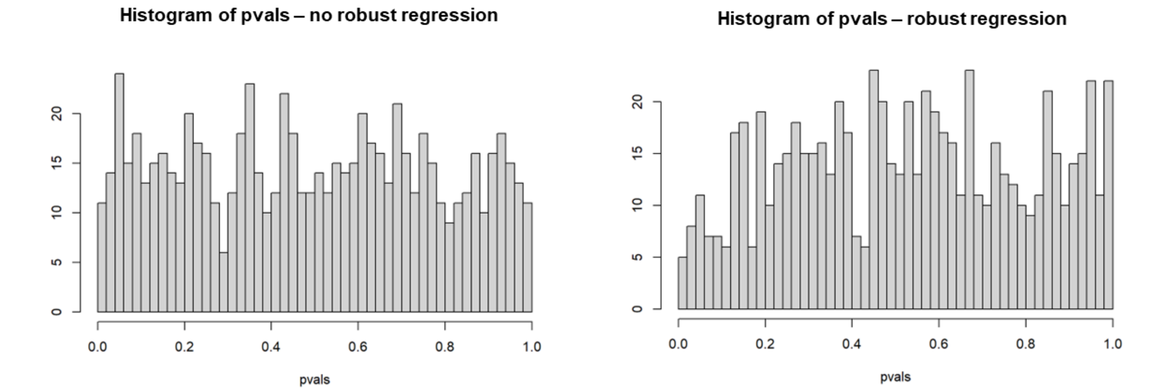

Peptidoform level

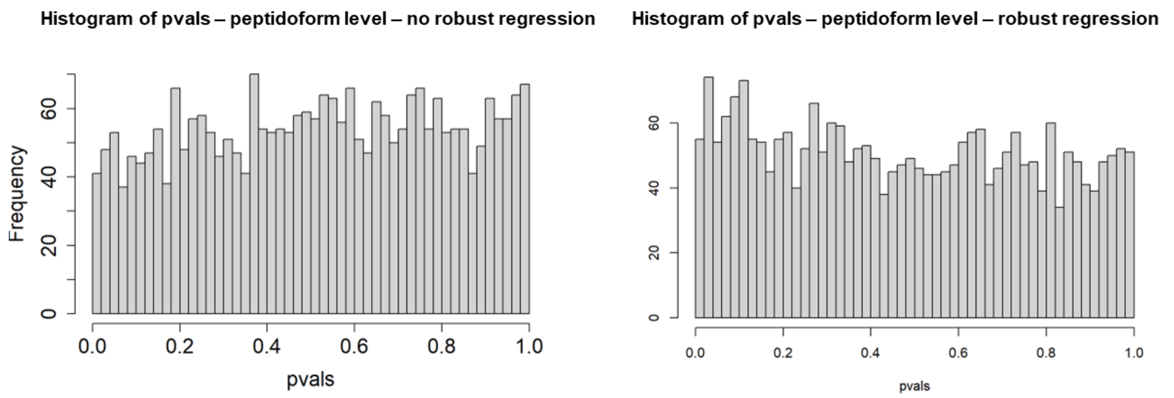

**Supplementary figure 12:** Distribution of p-values for the mock analysis of the phospho dataset including the use of a global profiling run, for analysis on PTM level (top) as well as peptidoform level (bottom). For the plots on the left, the analysis was carried out without robust regression during the modelling step. In contrast, the plots on the right correspond to an analysis carried out with robust regression during the modelling step.

*Example 3*

PTM level

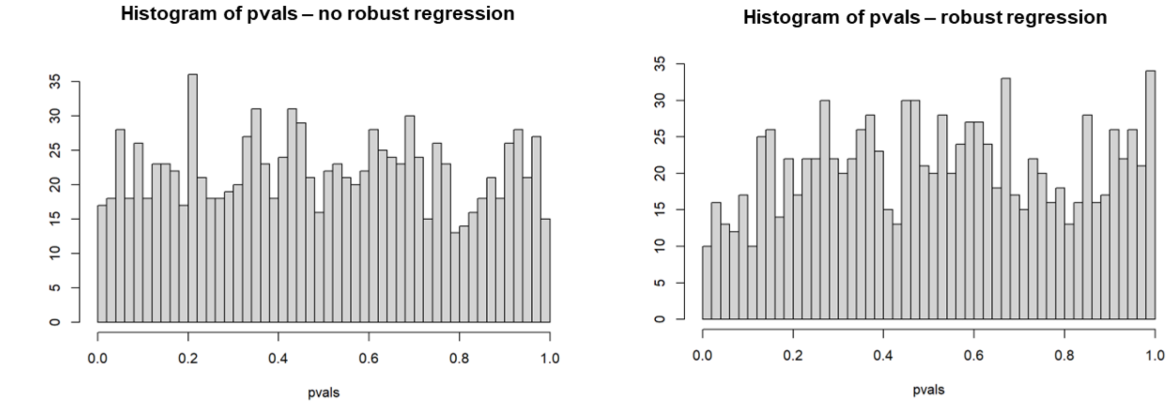

Peptidoform level

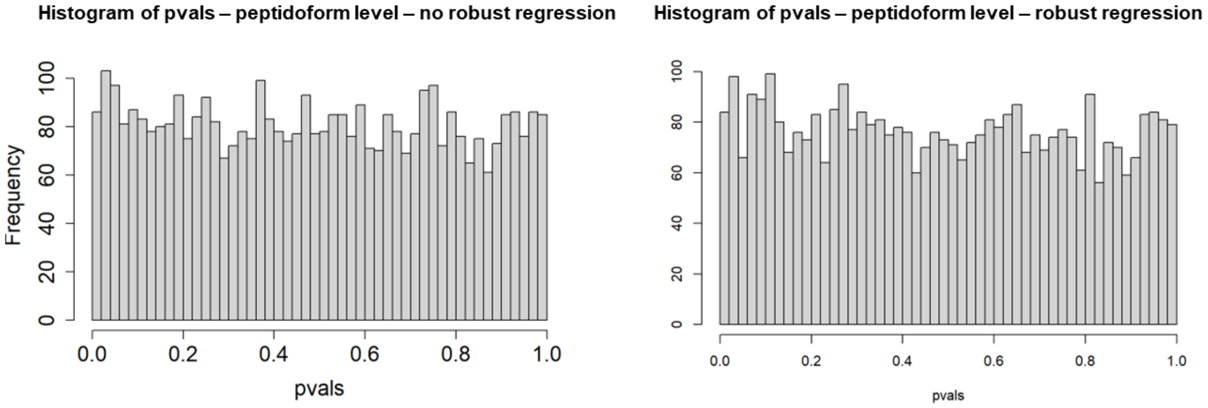

**Supplementary figure 13:** Distribution of p-values for the mock analysis of the phospho dataset including the use of a global profiling run, for analysis on PTM level (top) as well as peptidoform level (bottom). For the plots on the left, the analysis was carried out without robust regression during the modelling step. In contrast, the plots on the right correspond to an analysis carried out with robust regression during the modelling step.

*Example 4*

PTM level

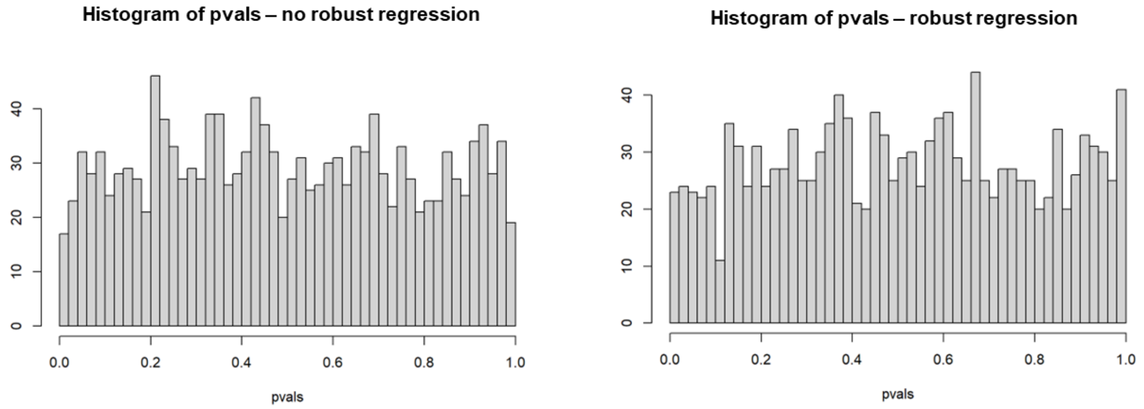

Peptidoform level

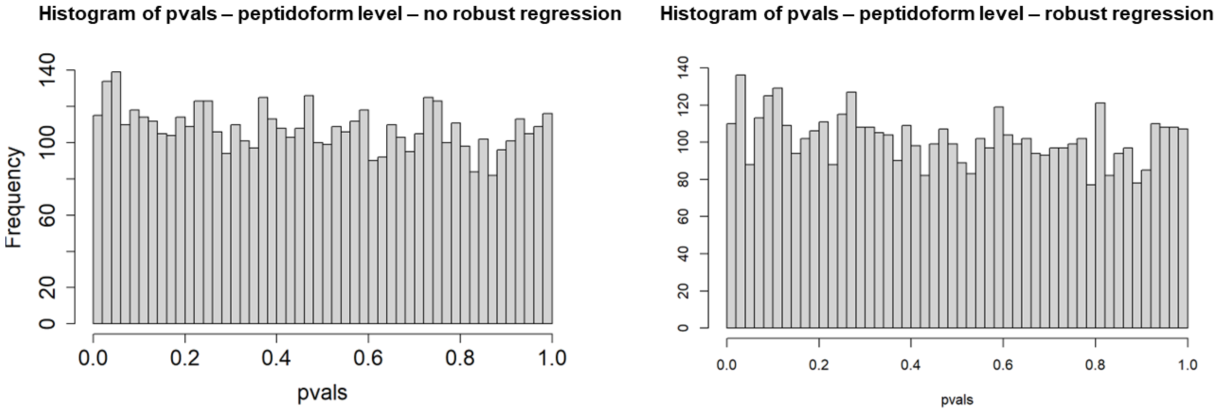

**Supplementary figure 14:** Distribution of p-values for the mock analysis of the phospho dataset including the use of a global profiling run, for analysis on PTM level (top) as well as peptidoform level (bottom). For the plots on the left, the analysis was carried out without robust regression during the modelling step. In contrast, the plots on the right correspond to an analysis carried out with robust regression during the modelling step.
